# Wireless Ensembles of Sub-mm Microimplants Communicating as a Network near 1 GHz in a Neural Application

**DOI:** 10.1101/2020.09.11.293829

**Authors:** Jihun Lee, Vincent Leung, Ah-Hyoung Lee, Jiannan Huang, Peter Asbeck, Patrick P. Mercier, Stephen Shellhammer, Lawrence Larson, Farah Laiwalla, Arto Nurmikko

## Abstract

Multichannel electrophysiological sensors and stimulators, especially those used for studying the nervous system, are most commonly based on monolithic microelectrode arrays. Such architecture limits the spatial flexibility of individual electrode placement, posing constraints for scaling to a large number of nodes, particularly across non-contiguous locations. We describe the design and fabrication of sub-millimeter size electronic microchips (“Neurograins”) which autonomously perform neural sensing or electrical microstimulation, with emphasis on their wireless networking and powering. An ∼1 GHz electromagnetic transcutaneous link to an external telecom hub enables bidirectional communication and control at the individual neurograin level. The link operates on a customized time division multiple access (TDMA) protocol designed to scale up to 1000 neurograins. The system is demonstrated as a cortical implant in a small animal (rat) model with anatomical limitations restricting the implant to 48 neurograins. We suggest that the neurograin approach can be generalized to overcome many scalability issues for wireless sensors and actuators as implantable microsystems.

## Introduction

Detecting and stimulating physiological electrical activity at multiple points helps to gain insight into the operation of targeted biological circuits such as those in the brain cortex. Current brain-machine cortical interfaces, for example, deploy monolithic microelectrode arrays (MEAs) to record multipoint signals from specific functional circuits such as the motor areas [1, 2, 3]. Elsewhere, mapping cardiac circuits by similar MEAs is pursued in diagnostic and therapeutic exploration [4, 5, 6]. Most high-performance neural sensors are based on micromachined silicon-based ‘beds of needles’ (MEA) or planar electrode arrays (ECoG) connected to external active electronics, with tissue contact points numbering on the scale of a hundred [7]. Scaling up the ‘channel number’ to many thousands is anticipated to provide a significant performance benefit, but doing so with current constructs presents challenges in architecture, data transfer, and implantability. One recent approach to scaling has employed microfabrication techniques to implement multiple multiplexed probe sites along rigid penetrating silicon shanks (e.g. Neuropixel approach [8]) and techniques for their high-throughput implantation are actively explored (e.g. Neuralink [9]). Wireless methods to eliminate percutaneous cabling to monolithic silicon sensors have also been reported (e.g. [10, 11, 12, 13, 14]).

A new approach to microscale sensors, proposed by several groups, envisions multichannel implants comprising ensembles of individual stand-alone microdevices. In one such approach, piezoelectric materials are integrated into the implant for energy harvesting and data transmission using ultrasound as the intermediary modality (‘neural dust’, [15]); neural signal recording and stimulation capabilities of these devices have been tested in vivo [16, 17]. Purely electromagnetic wireless power transfer (WPT) and communication in the near-field (inductive) have been studied e.g. by Ghovanloo and co-workers for mm-sized devices [18, 19, 20]. While achieving good electromagnetic coupling is inherently challenging for small area chip-scale antennas [15, 21], RF (Radio Frequency) techniques can, if properly optimized, provide an efficient path for wireless energy harvesting and telecommunication. We note other examples in which microdevices have demonstrated electrical microstimulation in small animals [22, 23, 24]. However, to our knowledge, what is missing from all prior work is the crucial aspect of networking ensembles of microimplants by a communication scheme that is scalable to the desired large numbers. We report in this paper the design and implementation of wirelessly networked ensembles of microdevices we term “neurograins”, with benchtop and in vivo demonstrations of neural recording and stimulation, respectively, in the rat cortex.

Figure 1 summarizes the salient features of the neurograin system where wireless communication-compatible microcircuits (ASICs) are realized in a 65 nm RF-CMOS process, each chiplet occupying a total volume of ∼0.1 mm^3^ (Fig. 1b). The electromagnetic interface for communication and near field power harvesting between an external hub and the ensemble of neurograins is conceptualized in Fig. 1a, which also shows the additional subcutaneous resonant co-planar relay coil designed to improve the WPT efficiency. Block diagram level description of the key electronic functions including the dedicated telemetry blocks are shown in Fig. 1c for the recording and stimulating versions of the neurograin system, respectively.

**Fig. 1.**
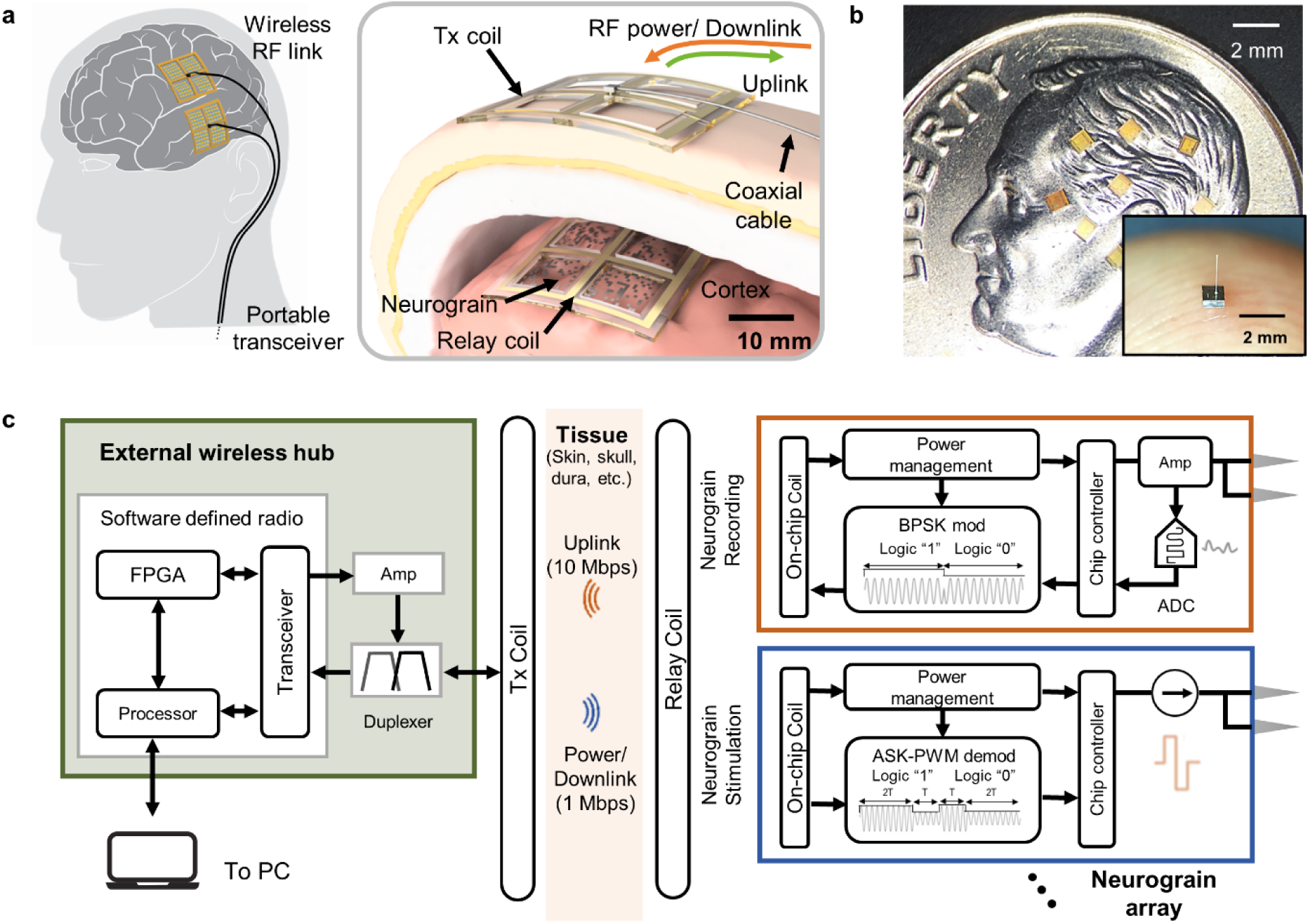
Wireless neurograin system for distributed autonomous networking. **a**, Concept of a transcutaneous RF power and data link for a neurograin array. **b**, Size scale of the present 650 μm × 650 μm × 250μm chiplets on a U.S. dime (inset: the optional post-process integrated microwire for intracortical access) **c**, Block diagram of the overall electronic system, consisting of the external RF hub, inductive coupling link, and key features of the neurograins for recording and stimulation, respectively.

Standard ASK, FSK, LSK, or IR-UWB modulation schemes have been used to communicate with microimplants, but demonstrated in limited channel bandwidth systems, typically below 1 Mbps [23, 25, 26]. For large ensembles of spatially distributed implants, a more versatile, and sophisticated networking protocol with high data bandwidth is needed. Multichannel techniques such as frequency-division multiple access (FDMA) and code-division multiple access (CDMA) are well-established in cellular networks [27]. However, when considering their adaptation to microimplants, the FDMA approaches encounter channel size restrictions imposed by the requirement for a high-Q microantenna [28]. CDMA in contrast requires complex microcircuitry, and has thus only been demonstrated as a two-channel system thus far [29].

In light of these considerations, we have opted to implement in our neural implants an alternative approach inspired by cellular and other multi-user networks — a specific power-efficient time-division multiple access (TDMA) network strategy for communication and active management of the system latency, which provides the necessary scalability and power and size efficiency. Supplementary Table 1 compares some state-of-the-art microdevices with the work presented here.

### System-on-Chip microimplant

Individual microchips operating as part of wirelessly networked ensembles need to incorporate multiple functions from energy harvesting to bidirectional data communication to neural recording and/or stimulation. The neurograins of Fig. 1 are sub-mm size systems-on-chip (SoC) designed as application-specific integrated circuits (ASIC) which must be tightly constrained for ultralow-power operation. In this work, we have designed ASICs using the TSMC 65 nm MS/RF LP CMOS foundry process, with no off-chip components. The microphotograph of Fig. 2a shows the overall footprint and main functional blocks of a neurograin chip fabricated for electrical microstimulation, in accordance with Fig. 2b. The corresponding block diagram and footprint for recording neurograin and others are shown in Supplementary Fig. 1.

**Fig. 2.**
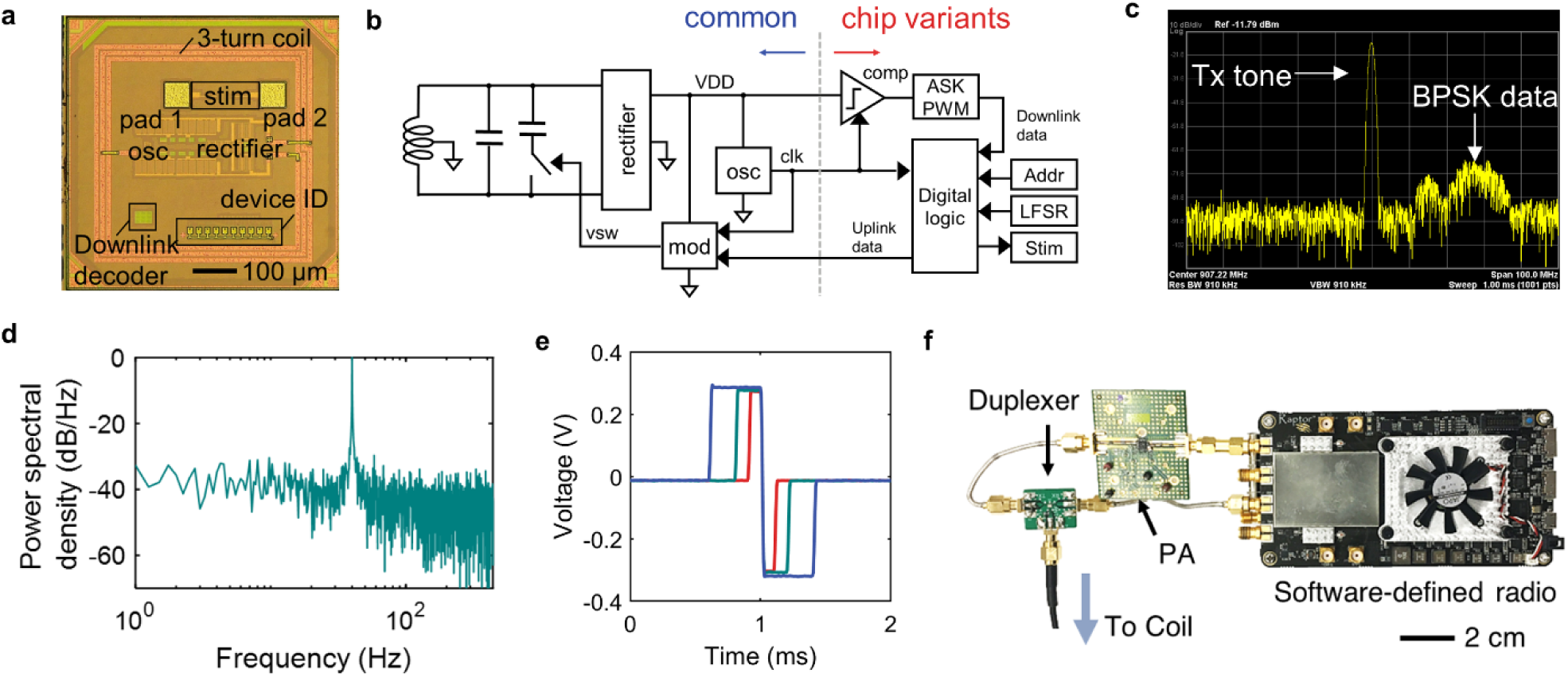
Design of neurograin chip and wireless power and data link. **a**, A photomicrograph of the stimulating neurograin with 500 μm diameter 3-turn microcoil, rectifier, downlink decoder and device ID. A pair of contact pads comprising the physiologic interface with tissue is also shown. **b**, ASIC circuit block diagram. The blue arrow highlights the RF interface (circuits common between all neurograins) while the red arrow points to blocks dedicated to particular physiologic interfaces (recording, stimulation, etc). **c**, RF spectrum showing the Tx tone and BPSK-modulated backscattered data lobe of the receiving channel. **d**, Normalized power spectral density (PSD) of a recording neurograin detecting a 40 Hz, 2 mV_ppk_ sinusoids injected into saline. **e**, Biphasic current stimulation voltage waveforms generated by the stimulating neurograin with three different pulse widths per phase (100, 200, 400 μs). A 20 kOhm load was used here to simulate tungsten electrode impedance at 1 kHz. **f**, Photograph of the benchtop external RF telecom hub consisting of software-defined radio (SDR), a power amplifier (PA), and a duplexer.

All neurograin chiplets share a common “RF interface” while executing unique and specific physiologic interfaces (common microcircuits indicated by the blue arrow in Fig. 2b, while the red arrow highlights one implementation of purpose-specific sub-circuit, for microstimulation in this case). This common interface notably includes an efficient RF power harvesting circuit, feeding off a three-turn on-chip coil (Fig. 2b), and utilizing a three-stage, low turn-on, deep n-well NMOS transistor rectifier circuit with low transistor threshold [30]. Low input voltage rectification, a critical feature for the feasibility of neurograin microimplants, is achieved through avoidance of body effects in the NMOS transistors deployed in this design. An RF backscattering data uplink channel at a rate of 10 Mbps operates near ∼1 GHz wireless powering link, and is enabled through a modulator (“mod” in Fig. 2b). The specific choice of wireless frequency near 1 GHz, based on electromagnetic simulations, represents our compromise between tradeoffs to achieve an acceptable near-field microantenna cross-section while minimizing the penalty by absorptive losses in body tissue.

We use a Binary Phase Shift Keying (BPSK) modulation for uplink communications. The modulator operates simply by toggling a tuning capacitor to enable RF backscatter while ensuring that the input impedance is maintained at a constant (maximized) magnitude through this phase modulation to guarantee a stable power supply [31]. Data is upconverted prior to backscatter using an on-chip, free-running relaxation oscillator with an operational frequency near 30 MHz; the latter enables spectral isolation of the BPSK-modulated signal from the transmitted RF powering (Tx) tone as shown in Fig. 2c. High fidelity isolation of the backscattered data facilitates low-noise demodulation and recovery at the external hub (Fig. 2f), where it is analyzed online or offline (see method). An example of the demodulation of the BPSK backscattered signal is shown in Supplementary Fig. 3a. In addition, the network architecture and data rates highlighted herein allow incorporation of downlink communication around the Tx tone to complete a real-time bidirectional communication loop as described in the next section. For neural stimulation applications, we choose a 1 Mbps downlink rate, sufficient to include command signals for commonly used electrical microstimulation protocols for the mammalian brain.

A particular challenge for designing a communication protocol for our ensembles of neurograins is the variance in the on-chip generated clock frequency and phase which arises from the variation in the power harvested by each chip. This variation makes synchronization difficult to achieve under normal protocols. We tackle this challenge by using a custom amplitude shift keying and pulse width modulation (ASK-PWM) scheme to implement the 1 Mbps downlink, as demonstrated by the circuit in Fig. 2b. This modulation approach provides network synchronicity and is robust across the measured range of chip clock rates. The ASK-PWM approach, which is compatible with our proposed networking architecture outlined in the next section, encodes each bit as a combination of high and low power (amplitude) pulses of variable duration. Received data on individual neurograins may thus be decoded using only the relative durations between the high and low states, independent of the internal clock frequency. Supplementary Fig. 2 illustrates the ASK-PWM scheme and the decoding circuit implemented in this work.

A unique device ID is imperative for the functional implementation of an addressable wireless network. For the neurograin microimplants, we have successfully tested two distinct ID approaches: (a) a physical digital address, engraved on-chip by laser ablation [32], and (b) an address scheme based on physically unclonable functions (PUF) [33, 34], discussed further below in a neural recording example. In the case of the laser ablation approach the use of which is demonstrated below for microstimulation (“device ID” in Fig. 2a), a user-selectable 10-bit address is physically imprinted onto an array of ten metal interconnect traces on the chip top aluminum metal layer by selectively ablating a specific number of traces with a 532 nm pulsed laser. The address circuits using the PUF approach take advantage of intrinsic 65 nm CMOS fab process variations to generate unique randomized bit patterns. The number of bits in a PUF address has to be carefully considered to minimize the chance for “collisions” due to address overlap for a given population of networked nodes. In our case, we have implemented 16 to 20-bit PUFs with the anticipation of an insignificant collision rate for ensembles of up to 1000 nodes. The PUF approach offers distinct advantages in scalability due to the compact footprint and lack of need for labor-intensive post-processing, although the unique address needs to be “discovered” for every neurograin.

The front end of a neurograin SoC integrates high-fidelity physiologic sensing (recording) and actuating (stimulation) interfaces, with a special focus on ultra-miniaturization and extremely low-power operation. To this end, we have designed a recording analog front end (AFE), a version of which is shown here for the acquisition of electrocorticographic (ECoG) signals using a DC-coupled V/I converter merged with an analog-to-digital converter (ADC) in an area-efficient capacitor-less approach [35]. The AFE directly senses neural potentials (without the need for bulky AC-coupling capacitors), and thus is able to occupy an overall footprint of 100 μm × 100 μm, while recording at a 1 kHz sampling rate and 8-bit resolution. Benchtop characterization measurements are summarized in Fig. 2d and Supplementary Fig. 3b.

Electrical microstimulation, on the other hand, is facilitated through circuits designed to deliver biphasic current pulses with programmable pulse widths (100, 200, and 400 μs), as commanded by the external wireless hub through the downlink (Fig. 2e,f). At present, the level of current stimulation depends on the chip power level and can range up to 25 μA as shown in Supplementary Fig. 3c; future work will incorporate stimulation amplitude regulation across the network. We implement programmable stimulation control to include both single shot as well as 100 Hz pulse train paradigms, the latter mitigating the data burden on the downlink. The recursive single-shot mode on the other hand requires significant communication bandwidth, but is capable of triggering burst stimulation at rates faster than 500 Hz, which is well above the frequency commonly used for electrical microstimulation.

### Networking of neurograin ensembles

A TDMA networking architecture enabling periodic, scheduled communication to and from ensembles of implanted devices is employed, using a single carrier frequency. The efficient use of finite channel bandwidth while maximizing the networked node count is of utmost importance in a neural prosthesis application, which also demands low latency data streaming capabilities.

The key tradeoff in large-scale network design is hardware complexity versus networking efficiency, the former defining stringent power and area constraints for the microdevices while the latter being critical for scalability. In recognition of these considerations, the neurograin SoC network was initially first realized using unidirectional “Autonomous” TDMA since it is a simple, compact and low-power approach for early ensemble testing at benchtop and in rodents, where channel counts are limited by anatomy. In this mode, there is no synchronization between the nodes. Each microdevice buffers and stores up to 100 ms of data (1000 bits), and this data is packetized and transmitted over 100 µs (10 Mbps rate) in a pseudo-randomly determined timeslot. This hardware-efficient approach, however, has network capacity for ∼100 nodes, beyond which the possibility of packet collisions (as shown in Supplementary Fig. 4a) increases in the free-floating microimplants due to unscheduled backscattering and chip clock frequency variations.

The implementation of an externally commanded, programmable, synchronized “call-and-respond” network offers a significant enhancement in both network efficiency and adaptive channel selectivity compared to “autonomous” operation, at the cost of integrating downlink communications on the microdevices. The current implementation of this architecture, integrated onto the neurograin SoCs, has yielded the capability to bidirectionally communicate with > 700 microdevices over a 100 ms period with > 70% bandwidth efficiency, and is theoretically extendable up to > 1000 microdevices through the integration of network pre-calibration for shorter “call” downlink (in ongoing design updates), thus being the ultimate candidate of choice for large-scale multi-node communication for distributed microsensor networks.

The “autonomous TDMA” protocol relies on the chip’s unique device ID, and the chip is in “transmit-only” operation. This prefabricated PUF ID probabilistically determines the device’s initial communication time-slot in the network queue. Data is then periodically backscattered at the predesigned network latency (nominally 100 ms). Microdevice on-chip clocks are designed to be free-running at 30 MHz with +/-12.3%; this variation is due to the spatial dependence of the magnetic coupling strength and resulting varied received RF power. Fig. 3b shows the measured transient data received at the external hub from a network of 64 chips. The “pulses” indicate periodic data packets sent by the nodes. For each data packet, decoded bits are compared with the predefined sequence to determine the bit-error-rate (BER). We recovered data with an average BER of 0.007% from 61 data packets detected during a 100 ms interval, while 4 additional packets were corrupted due to packet collisions showing BER >1% (Supplementary Fig. 4b). Characterization of the long-term stability of the WPT and data telemetry scheme is demonstrated in Fig. 4c. When we subsampled 1800 data packets from 64 neurograins over 30 minutes, the spread of their relative pulse powers demonstrated the variable energy harvested as a function of the chip’s spatial location. Packet RF power and recovered chip clock frequency are plotted, with each color representing an individual neurograin, and k-mean clustering into 64 groups shows a well-isolated, stable clock frequency over time.

**Fig. 3.**
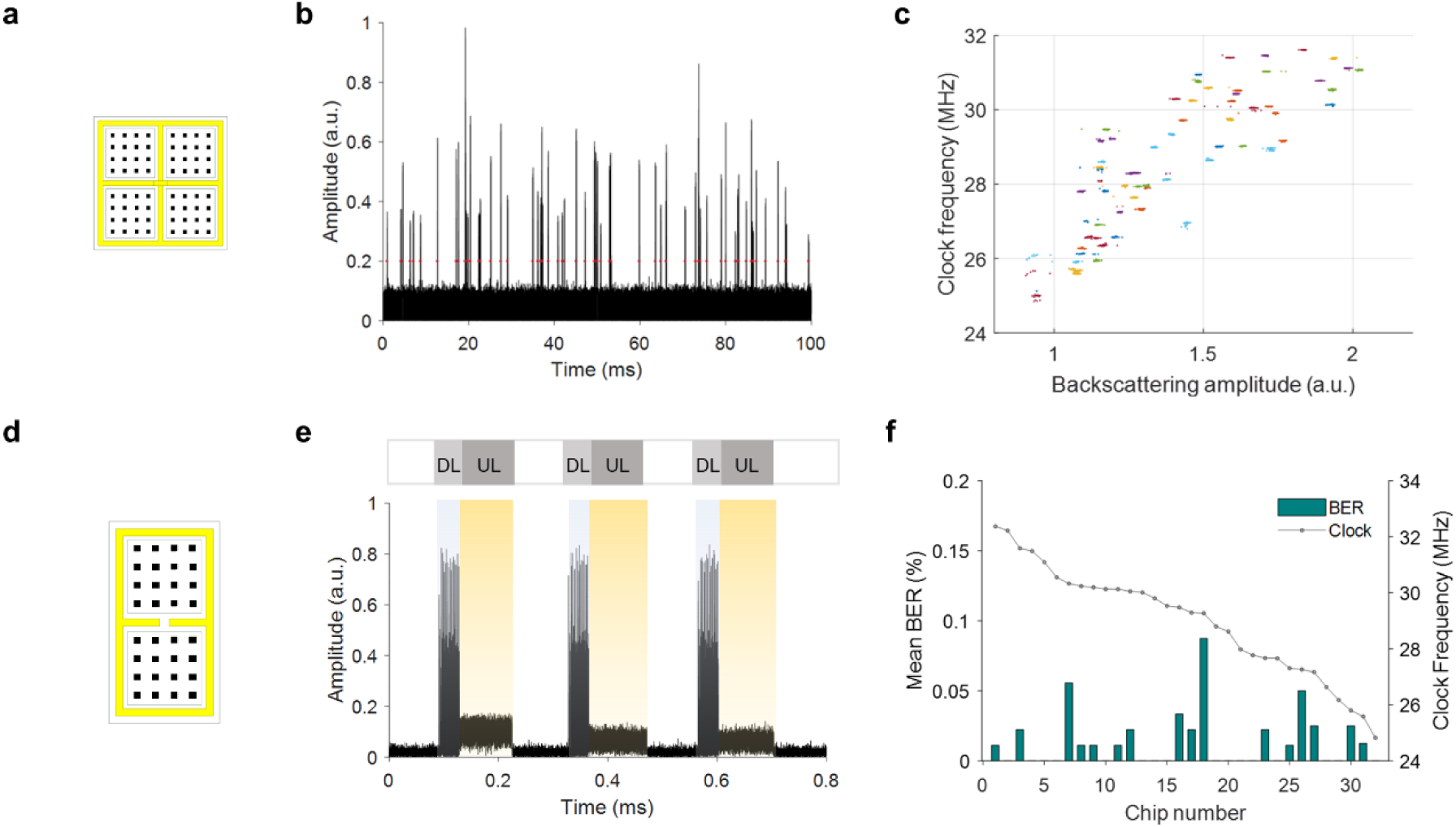
Data communication demonstration. **a**, 64 autonomous TDMA chips, with placement within a relay coil. **b**, A transient recording on uplink data packets from multiple chips under autonomous TDMA. **c**, Recovered clock frequencies and backscattering amplitudes of 1800 data packets sampled for over 30 minutes from 64 chips. **d**, A 32-node call-and-respond network within a relay coil. **e**, A transient recording on the downlink (DL) and uplink (UL) under call-and-respond TDMA. **f**, Averaged BER from 1000 uplinked bits versus chip clock frequency of 32 chips individually identified by PUF addresses.

**Fig. 4.**
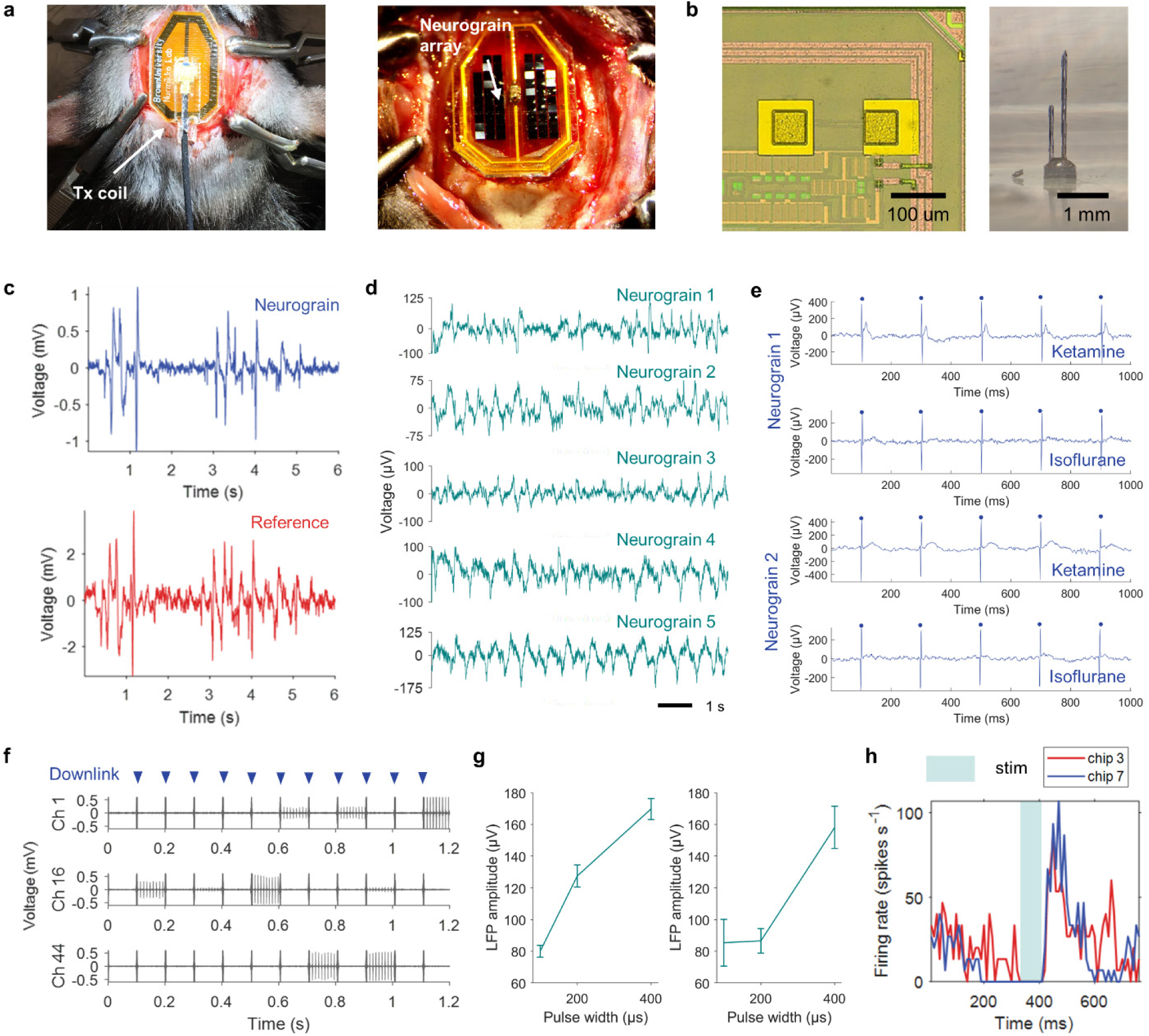
Neurograin recording and stimulation in saline and in vivo. **a**, A photograph of in-vivo implant and head-mounted components. **b**, photomicrographs on gold electrodes fabricated on-chip (left) and microstimulator with intra-cortical tungsten electrodes (right). **c**, Injected ECoG signal recordings from neurograin (top) and commercial device (Ripple neuro nomad) (bottom) in the saline. **d**, Multi-channel neurograin recordings on a spontaneous low-frequency oscillatory wave from the anesthetized rat. **e**, Recorded raw signals on 5 Hz electrical stimulation (blue dots) and evoked neural response under ketamine and 3% isoflurane using recording neurograin array. **f**, 100 Hz electrical stimulation trains from stimulation neurograin array in saline with downlink artifacts shown by blue arrows. **g**, Amplitudes of post-stimulus LFP response depending on the stimulation pulse width (per phase) from chip 3 (left) and chip 7 (right). Up to 25 μA current was delivered to the tissue. The error bar represents the standard deviation. **h**, The sum of the firing rate from 3 recording channels before and after the 400 Hz burst stimulation from neurograin (averaged over 5 trials and the bin size is 10 ms) which shows increased post-stimulus firing rates.

The “call-and-respond” TDMA network, in contrast, integrates the ASK-PWM protocol for downlink communication to the microdevices. The unique device identifiers (16-b PUF address), implemented on-chip, are recovered from the microdevices through an automatic start-up communication. These IDs are subsequently used to externally command individual devices through the ASK-PWM link. Each downlink command is received and downlink codes are deciphered across the ensemble in a synchronized manner. If the received downlink code matches a given device’s address, it transmits a data packet (“respond”). Fig. 3e shows the snapshot of the transient received signal at the external hub with the leaked, strong downlink call of 40 μs and the weak, uplink backscattered data of 100 μs for a network of 32 neurograins. Successful communications, measured by response rate to the downlink, occur with an average efficacy of 99% across this network, but we are also able to identify 3 other devices with <90% efficacy, possibly due to CMOS process variations (Supplementary Fig. 5a). The uplinked data from these chips, comprising 1000-bit predefined sequences, yielded an average BER of 0.013%, as shown in Fig. 3f (Supplementary Fig. 5b for each packet) validating the ability of the downlink/ uplink and the network to withstand variations in clock frequency and on-chip power.

### Wireless recording and microstimulation by neurograins

We have built and assessed the fully functional system of Fig. 1, both on the benchtop (saline solution) and in the in-vivo rat model for both neural recording and electrical microstimulation. For multichannel neural recording from ensembles of neurograins, the external wireless hub receives neural data as TDMA digital streams. In the stimulation mode, the hub coordinates the ASK-PWM downlink for space-time specific microstimulation.

Converting fabricated microchips into functional microsensors/stimulators requires further post-processing after their dicing from the wafer. For a fast turn-around, we have developed a polymer-based microfabrication technique [32] to deposit planar gold microelectrodes (60 μm pad size) atop the chip aluminum input contact pads. This approach is able to achieve electrode impedances suitable for epicortical ECoG recording (Fig. 4b left), and is suitable for attachment of microwires (a 50 μm diameter tungsten wire in Fig. 4b right) for use as intracortical electrodes such as used in the microstimulation experiments. For long term chronic device implantation, not the goal of this paper, we have reported elsewhere the development of ultrathin dielectric coatings, applied by atomic vapor deposition, to obtain biocompatible, hermetically sealed neurograins [36].

While each modality (recording, stimulation) is anticipated to ultimately integrate a single, bidirectional TDMA link, we are currently able to leverage this protocol to fulfill the specific needs of each type of neurograin ensemble. The recording chip reported here generates the uplink under autonomous TDMA (see the previous section). Since the head/skull size of a rat limits the number of practically implantable neurograins (here N=48), the probability of data packet collision is low. We are able to communicate with such ensembles while identifying the 48 unique PUF addresses (see Supplementary section on measured PUF performances and results in Supplementary Fig. 6). The unique address on each chip ensures continuously identifiable neural data flow from each recording neurograin. As examples of recordings in saline (Supplementary Fig. 7a), various current waveforms are injected from a nearby microwire either as proxy neural signals (Fig. 4c) or sinusoids at 200 and 40 Hz (Supplementary Fig. 7b,c) and the recordings by the neurograins are compared with those acquired by a commercial wired neural recording system (the multichannel neural signal processor “Nomad” by Ripple Inc).

For testing the neurograin system in the in-vivo rat model (acute), we used a 6-channel commercial intracortical stimulation device using tungsten microwire electrodes while the epicortical neurograins recorded the evoked neural activity. The stimulation pulse width was set at 1 millisecond per phase and current up to 50 μA was delivered. An array of recording neurograins was placed on the animal’s cortex, covering most of the motor and sensory areas in proximity to the implanted wired stimulation device, and the external transmission coil (Tx) was positioned above the animal’s head (Fig. 4a). As shown in Fig. 4d and Supplementary Fig. 8a,b, in the absence of external stimulus, the neurograin devices were able to record spontaneous ECoG signals, prominently capturing the sub-Hz slow wave oscillations characteristic of Ketamine anesthesia [37]. In contrast, when the cortex was actively stimulated using 5 Hz current pulses, the neurograin ensemble captured clear post-stimulus evoked field responses (irrespective of the stimulation artifacts). The results of raw recordings are shown in the data of Fig. 4e. Superimposed traces in Supplementary Fig. 8c from multiple trials show that the recorded evoked responses persist for 20 to 70 milliseconds after a stimulation event. In contrast, recorded responses under isoflurane anesthesia (which inhibits cortical activity), conspicuously lack these evoked potentials, thereby eliminating the likelihood of their origin from spurious internal noise or stimulus artifacts.

In order to assess the stimulating-type neurograins, we demonstrate here the laser inscribed address approach, where we schedule the delivery of electrical stimulation parameters for each target chip through the use of the TDMA ASK-PWM downlink. Fig. 4f demonstrates programmed, patterned delivery of electrical stimulation pulses across a neurograin array; RF modulation by the downlink communication manifests as an ‘artifact’ (used here for timing reference). We have verified that ensembles of chips are individually responsive to the downlink commands, and able to deliver electrical microstimulation in a spatially distributed manner. Supplementary Fig. 9a shows downlink coordinated stimulation trains from a population of 12 neurograins, each following reliably its downlink command, here over 3 cycles.

For acute rat experiments with implanted ensembles of stimulating neurograins, we microfabricated the auxiliary device of Supplementary Fig. 9b where an array of conventional recording microwires was integrated into the same polymer platform as the wireless stimulating microdevices, and connected to commercial multichannel (wired) recording electronics. This hybrid device enabled detailed validation of the physiologically relevant neural activation upon the operation of stimulation neurograin. As one example, we targeted a single neurograin in the implanted ensemble, and programmed it to inject a 100 Hz intracortical stimulus pulse train (up to 25 μA current). We were able to record post-stimulus LFP responses from multiple recording channels (Supplementary Fig. 10a). This stimulation induced 100 Hz oscillatory neural activity appeared to be widespread across the cortex, indicating that a single stimulation target may be able to modulate wide cortical areas. The amplitudes of the evoked responses showed the well-known dependency (in electrophysiology literature [38, 39]) on stimulus pulse widths, whereby longer pulses evoked the largest LFP responses (Fig. 4g). In another neuromodulation example, we delivered fast 400 Hz stimulation pulse trains in 100 ms blocks, and were able to observe the characteristic neuronal bursting with increased post-stimulus firing rates described in literature [37, 40] under ketamine anesthesia (demonstrated in Fig. 4h and Supplementary Fig. 10b).

### Characterizing Wireless Power Transfer for Optimal Performance

To optimize the efficiency of electromagnetic data and power transfer between the external hub and ensembles of spatially distributed neurograin implants, we have utilized a three-coil architecture for inductive coupling. Design constraints for this coil system, comprising the external Tx coil, the implanted Tx-size-matched relay coil and the on-chip receiver (Rx) microcoils, are informed by high-performance electromagnetic simulations. The Tx and relay coils are both designed in a geometry which can be viewed as a *de facto* superposition of unit sub-coils, capable of linear extension for enhanced areal coverage while maintaining WPT efficiency as shown in Fig. 5a and Supplementary Fig 11a,c. Leveraging the same underlying electromagnetic principles, we have been able to custom design Tx/Relay coil geometries to tailor different versions of the system to optimally accommodate the anatomical structure of both future primate experiments and the present rodent model (dimensions and simulation parameters shown in Supplementary Table 2). In each case, the characteristic coil layout generates current flow patterns (Supplementary Fig. 11b,d) whereby the respective directions of magnetic flux of adjacent subunit coils are opposite to each other as in Fig. 5b. The design permits lateral expansion of the power harvesting area, without deteriorating coil quality factors or coupling factors between coils. The on-chip microantenna design (Rx) is optimized as a 3-turn Rx coil using the thick aluminum top metal layer available within our semiconductor foundry process. The coil has an outer diameter of 500 μm, and is able to accumulate magnetic flux with an efficiency compatible with powering on-chip operations of the high-load/low-power ASIC at 1 GHz (Fig. 2a).

**Fig. 5.**
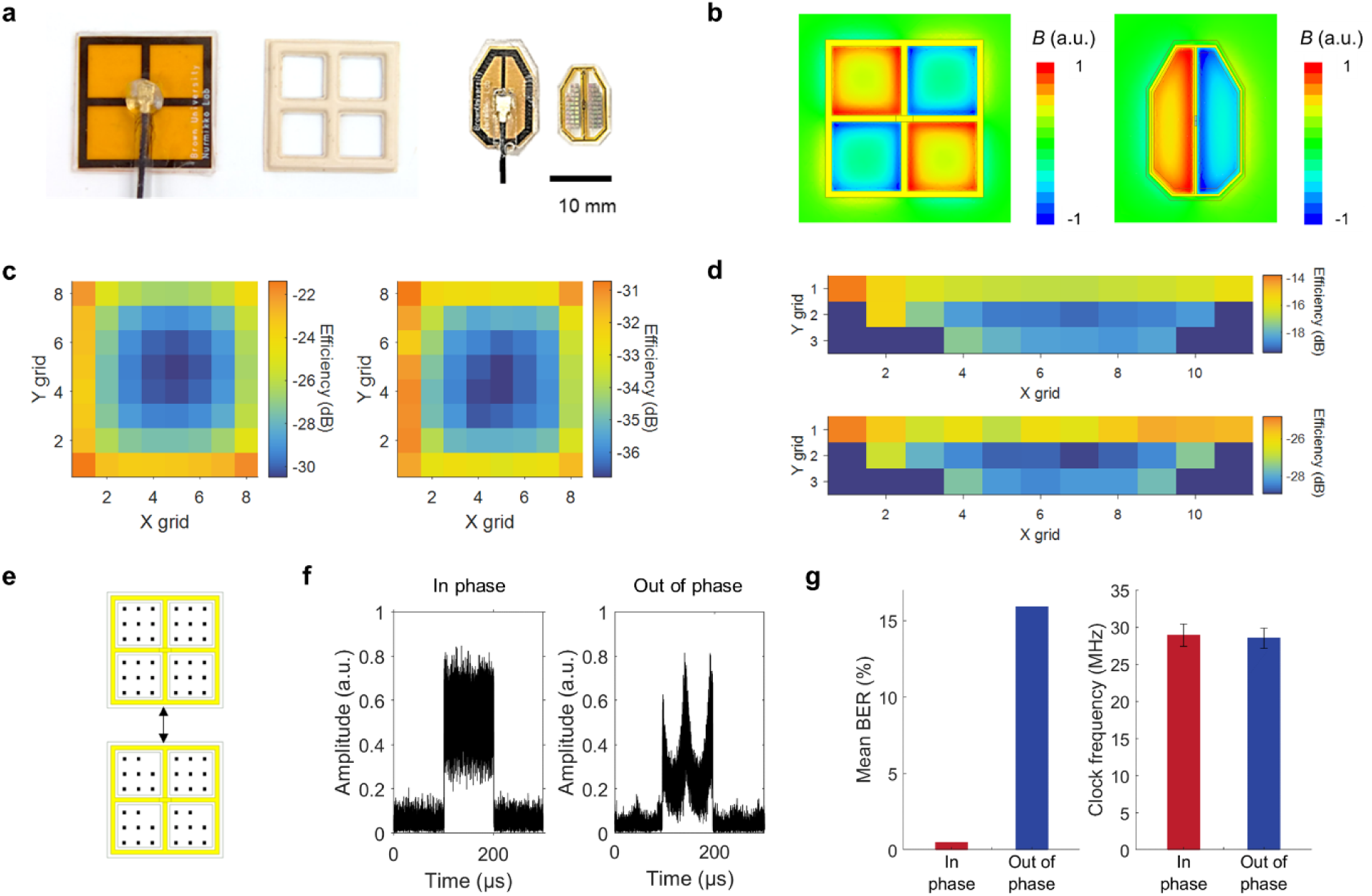
Wireless efficiency characterization. **a**, Photographs of packaged Tx and relay coil for the primate and rodent model with embedded chip array. **b**, The magnetic flux generated by the coil design which shows alternating fluxes of subunit coils. (the direction of the flux is shown as red or blue) **c-d**, Maps of simulated (left, top) and measured (right, bottom) wireless transfer efficiency regards to the spatial location inside one of the subunit relay coil for the primate and rodent model, respectively. X grid size is 1 mm for both cases and y gird size is 1 mm for the primate model coil and 0.8 mm for the rodent model coil. **e**, Parallel-relay scheme with 69 chips activated through two adjacent relay coils with 8 mm separation demonstrating the scalability of wireless powering and data communication area. **f**, Received RF data packet shapes using two RF sources in-phase (left) and out of phase (right). **g**, Averaged BER and clock frequency of 865 and 733 data packets (in-phase and out of phase, respectively).

One way to quantitatively analyze the electromagnetics of the wireless link *in-situ* is to measure the clock frequency of the chip, which serves as an estimate of the power level received by the chip, as per the simulated frequency-power relationship in Supplementary Fig. 3d. We have used a liquid-form head phantom construct as a proxy for a physiologically relevant RF environment, and analyzed the effect of tissue path loss and surrounding tissue permittivity on the WPT and data link [41]. We have mapped the WPT efficiency at 64 different spatial positions inside a primate model relay coil using a location sampling grid with an x-y pitch of 1 mm (Fig. 5c), and in a rodent model coil at 26 positions with an x-y pitch of 1 mm and 0.8 mm, respectively (Fig. 5d). Based on the consideration of anatomical characteristics, the separation between the Tx coil to relay coil was 8 and 5 mm in the primate and rodent model coil, respectively.

As expected, the Rx coil achieves its highest efficiency when in close spatial proximity to the relay coil perimeter, and coupling decreases when moving toward the center of each subunit coil (Fig. 5c,d). For the primate model coil, the WPT efficiency ranges from −36.77 dB to −30.73 dB (center to edge). In the rodent model coil, the maximum efficiency of −24.88 dB is located at the corners decreasing to −28.90 dB at the center. Received power at the microimplant also depends on the horizontal and angular alignment of Tx coils as shown in Supplementary Fig. 11e, f. For the primate model, lateral movement (x-y plane) of the Tx coil results in a loss of 1.3 dB for a 3 mm horizontal displacement. For the asymmetric coil morphology used in the in-vivo rodent experiments, a y-directional misalignment of 3 mm resulted in a 0.6 dB penalty, while an x-directional misalignment resulted in a much larger 6.73 dB loss, which is expected given the coil aspect ratios. A 20-degree tilt resulted in 1.29 dB vs 1.9 dB loss in WPT efficiency in the primate and rodent configurations, respectively. In the actual operation of the neurograin system, we used the strength of the backscattered RF signal as the guide to optimally align the Tx coil with submillimeter accuracy.

This wireless powering and data transmission scheme is intrinsically designed to be parallel-extendable to cover multiple cortical areas and increase channel capacity. As an example of this scalability, Fig. 5e demonstrates a dual coil and dual Rx channel, 69-node system using autonomous TDMA chips. When the dual Tx is in phase, implemented simply through sharing an RF source in our WPT design, each subunit coil drives current in a direction opposite to that of adjacent coils. Therefore, their respective magnetic fluxes do not destructively co-interfere, as depicted by the retained packet shapes in Fig. 5f-left, and the averaged BER is 0.5% over 865 packets. When the two Tx coils are out of phase, which is the common case for two regular coils, destructive interference of their magnetic fields results in deformed packet shapes (Fig. 5f-right). The mean BER over 733 data packets rises to 15.9%. Packet collisions are excluded for both configurations. Average clock frequencies across chips are similar in both condition (Fig. 5g-right plot), which suggests that the dual-channel system, by only using a common RF source, can deliver wireless energy to the larger number of spatially dispersed nodes without risk of significant cross-talk or BER deterioration.

### Assessment of specific absorption rate of wireless power transfer

In consideration of the effects of radiation on tissue, Supplementary Fig. 12 shows computed specific absorption rate (SAR) maps [41]. According to present IEEE guidance (and related FDA dictum), adverse effects are not expected below the 10 W/kg SAR ‘threshold’ value [42]. Applying this standard, our primate coil system can transfer enough power for current ASICs to operate to 48.4% of the lateral relay coil area which can include approximately 500 neurograins (Supplementary Fig. 12a). For the rodent coil and values simulated for our experiments, an input power of 0.072 W (18.6 dBm) would provide sufficient RF energy to neurograins across the relay coil, at the equivalent peak spatial SAR of 3.69 W/kg (Supplementary Fig. 12b). Actual measured WPT efficiency was, on average, 7.4 dB and 7.7 dB lower than the simulated values in primate and rat model coil, respectively, although the spatial dependence of the measured efficiency generally followed the simulated results. We note that the power measurement includes losses at multiple stages across the link from the RF source to the on-chip circuit. The overall efficiency can be further increased, for example, by improving the on-chip rectifier efficiency, or better tailoring the impedance match in the coil/RF circuitry. Due to the specifics of the TSMC 65nm foundry process, the embedded planarization-enabling filling structures across the multiple metal layers of the ASIC chip may have further limited the cross-section and coupling factor of the Rx coil. Further improvement on peak SAR will be dependent on identifying the sources of the reported additional 7 dB measured loss in our system and mitigating these causes. When achieving the simulated maximal efficiency, 0.1 W (20 dBm) of Tx power in the primate model coil can provide adequate energy to a neurograin ASIC at any location within the relay area, which corresponds to a peak spatial SAR of 3.66 W/kg, averaged over 10 g according to results.

Lastly, Supplementary Fig. 12c shows the effect of an extra adjacent Tx/relay coil on the SAR value, which shows that adding another RF WPT/data path (e.g. for cortical coverage for more than one area) did not affect the local SAR value. Thus, from the SAR point of view, our wireless system can scale up using multiple RF paths (such as for neurograin populations in the thousands) with minimal impact on the peak local SAR values, while distributing RF energy across the multiple cortical areas.

## Conclusions

We have described here what we believe is the first realization of a scalable implantable wireless microsensor and microstimulator network. Each sub-mm sized silicon chip (0.1 mm^3^), a neurograin, combines RF energy harvesting, data communication, and recording or stimulation capability as an autonomous unit with its unique identifier. We have designed a custom time-division multiplexing approach in building a spatially distributed microsensor system which communicates with an external wireless hub. The multichannel system has been demonstrated in-vivo rat model to record epicortical ECoG signals and intracortically stimulate neuronal microcircuits, respectively. While the head size of the rodent model has limited us to 48 implanted neurograins in this paper, the networking approach has the capability to be scale up to 1000 nodes. The electronic design principles of the neurograin SoC circuits can in principle be ported to further deep sub-micron CMOS process nodes (such as the 22nm node), reducing the chip volume by another order of magnitude. This can provide a pathway to minimally obtrusive implantation of large populations of devices such as by recently developed intracortical insertion techniques [43]. Importantly, our bidirectional wireless communication approach lends itself to real-time ‘adaptive sensing’, i.e. focusing on those sub-populations of neurograins which are most directly engaged in information exchange with the underlying biological circuits. In a broader sense, we may have demonstrated a step towards building versatile wireless electronic biointerfaces such as for future brain-machine interfaces, not unlike early mobile phones whose evolution to smartphones has transformed human communications.

## Methods

### Equivalent circuit design

In a 2-coil system shown in Supplementary Fig. 13a, the efficiency of wireless power delivery (η) between the Tx coil and Rx coil can be described by

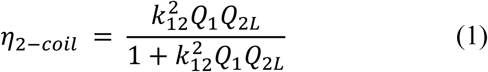

where *k*_12_ is the coupling factor between the Tx and Rx coil and *Q*_1_ = *ω*_0_*L*_1_/*R*_1_ and *Q*_2*L*_ = *Q*_2_/(*Q*_2_*Q*_*L*_ + 1), where *Q*_2_ = *ω*_0_*L*_2_/*R*_2_ and *Q*_*L*_ = *R*_*L*_/*ω*_0_*L*_2_ [44, 45]. Here, due to load transform from the resonant capacitor *C*_2_, *R*_*L*_ can be solved as 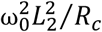 and *R*_*c*_ is the circuit equivalent impedance [28]. *k*_12_ depends on the size of the coils and the distance and arrangement between two coils. In the case of on-chip micro-coils implanted, e.g., on the surface of the cortex, the size mismatch between Tx and Rx antennas is significant (where the latter are ∼0.25 mm^2^ in our system) so that, given the separation between two coils exceeding several millimeters, the maximal coupling *k*_12_ is severely limited.

In the 3-coil system, a secondary coil can be introduced so that it can have moderate coupling both with Tx and Rx coils, and thus increases the overall efficiency of WPT (Supplementary Fig. 13b). The coupling factor between the relay and Rx coil, *k*_*R*2_, can be ten times larger than *k*_12_ in the co-planar arrangement and the efficiency in 3-coil wireless powering can be simply written as

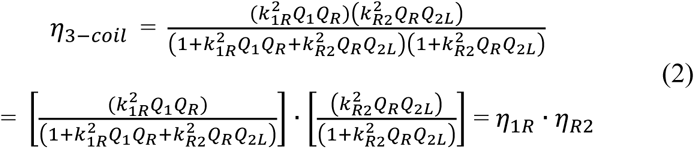

where *k*_1*R*_ is the coupling factor between the Tx and the relay coil, and *k*_*R*2_ is the coupling factor between the relay and Rx coil [44]. *Q*_*R*_ is the unloaded quality factor of the relay coil. Using a secondary coil that has a size comparable to the Tx coil, *k*_1*R*_ can be increased significantly, and a high *η*_1*R*_ can be achieved. Also, co-planar placement of the relay and Rx coils can improve *k*_*RR*2_, which results in higher *η*_*R*2_. Note that since the relay coil is not directly loaded by source or load impedance, *η*_1*R*_ and *η*_*R*2_ benefits from a high-quality factor for the relay coil *Q*_*R*_. Consequently, the relay coil collects high magnetic energy to the target area, improving power transfer efficiency for the ensemble sub-mm Rx coils. Based on simulated coil parameters in Supplementary Table 2 and equation (1), (2), the maximum and minimum efficiency within the relay coil are calculated in the primate and rodent model coil for both cases of 2 coil and 3 coil as shown in Supplementary Fig. 13c. The calculated efficiency complies with the 3-coil simulation in Fig. 5c,d and also reveals more than 15 dB efficiency increases in the 3-coil system compared to 2-coil.

### Electromagnetic simulation

The design of coils was performed and simulated on a High-Frequency structure simulator (HFSS ANSYS electromagnetic suite, Ansys) with tissue models including their dielectric properties around 1 GHz. Design parameters on the Tx and relay coil were chosen to maximize WPT efficiency within the power projection area while the maximum area is limited by the self-resonance of coils. In the rodent WPT system, the size of the coil was limited by the size of the rat skull. The On-chip Rx coil was also designed on electromagnetic simulation to improve WPT efficiency and the impedance matching with the high-load/ low-power ASIC. S-parameters from HFSS simulation results were imported on Cadence Design Systems Mixed-Signal Simulator to set up a full RF simulation test bench for wireless circuit design.

### Chip post-processing and packaging

60 μm sized pads on bare ASIC offer electrical path to biological tissue while the aluminum surface of them is not suitable for long-term tissue interface. For recording neurograin, the 60 μm × 60 μm gold electrode surface can provide appropriate impedance (300 kOhm at 1 kHz) to meet the input requirement of capacitor-less AFE for ECoG recording. And tungsten penetrating electrodes have an impedance low enough for stimulation neurograin, which has a nominal 10 kOhm impedance at 1 kHz. Using the polymer-based customized technique and photolithography [32], we have deposited 50 nm titanium and 200 nm gold pattern directly on the pad to serve as electrodes. For the stimulation chip, 1.5 mm-long tungsten wires with 50 μm diameter were manually attached to pads with silver epoxy (EPO-TEK® h20e) and coved with epoxy resin (Scotch-Weld™ DP100). After the electrode fabrication, implants were encapsulated by parylene-C except for electrical contacts for recording or stimulation to achieve the long-term biocompatibility.

### Coil fabrication, assembly, and packaging

The Tx and relay coil were fabricated on a polyimide substrate with a 0.1 mm thickness under a 2-layer PCB process using a 2 oz copper trace. For the Tx coil, trace width up to 1 mm was used to improve the quality factor while, in the relay coil, the trace width of 0.2 mm is used to reduce the effect of surrounding dielectric tissue. These design parameters were optimized in electromagnetic simulation before the fabrication. Coils were then assembled with matching capacitors and encapsulated with PDMS (Dow® SYLGARD™ 182) or 3M biomedical-grade epoxy resin. For Tx coils, 50 ohm mini-RF coaxial connector and cable are also attached. PDMS packaging processing has been done with an acrylic mold. Before the packaging, coils were cleaned with isopropyl alcohol, acetone, and DI water, and the mold was coated with anti-adhesion spray. The base PDMS layer with 0.5 mm thickness was cured for 3 hours at 100 °C, and attached to polyimide coils. The top PDMS layer covered again, resulting in an overall packaging thickness of 1 mm. Microimplant chips are tested without the package or packaged in PDMS or parylene-C. In the cases that longer-term hermetic packaging is needed, the relay coils are encapsulated with liquid crystal polymer by using a metal jig and a press machine [46].

### Data telemetry and demodulation

SDR (Raptor, Rincon Research Corporation or FMCOMMS3, Analog Device) consisting of an Analog Devices AD9361 RF transceiver and a Zynq SoC was used to generate RF transmission energy and to communicate with ASICs [47]. The SDR generates transmitting tone at 915 MHz, which is further amplified with an IC-based power amplifier (ADL5605-EVALZ, Analog Devices) board. An RF SAW duplexer (D5DA942M5K2S2, Taiyo Yuden) isolates transmitting and receiving paths. The SDR receiver front-end performs signal amplification, down conversion (945 MHz), and digitization at 30-45 MSa/s. The digitized IQ data was then ported to a PC for offline processing (in MATLAB/Simulink), which can be embedded into Zynq SoC for the onboard demodulation. The digital bitstream is decoded from the digitized IQ data after filtering, automatic gain control, coarse and fine frequency compensation, bit strobe correction, and BPSK demodulation. These processes have been optimized for the neurograin communication scheme. Automatic gain control normalizes the uplink data packet power which varies between chips. Frequency compensation and bit strobe correction processes, recovering the clock frequency of each chip, compensates the clock variance between chips to improve demodulation performance.

### RF wireless experiment setup

Liquid head phantom with the permittivity of 41.8 and the conductivity of 0.97 S/m around 900 MHz was made in accordance with the IEEE standard 1528-2013 to mimic the dielectric properties of head tissue [41]. The liquid phantom consisted of 64.81% 1,2-Propanediol, 0.79% NaCl and 34.4% DI water. A phantom bath was designed for the wireless test using acrylic plates and a 0.1 mm thin plastic sheet to reduce RF interference (Supplementary Fig. 14). The separation between Tx and relay coils was 8 mm or 5 mm and the gap was filled with the liquid phantom. Two plastic sheets supported the liquid phantom and the relay coil, respectively. For the efficiency measurement experiment, grid paper was attached inside the relay coil to guide the manual placement of the neurograin chip. For the Tx alignment, a 3D-printed Tx coil holder was placed below the bath, and above the paper with grid marks which provides references for the alignment between the Tx and relay coil.

### In vivo experiment

A wild-type Long-Evans adult male rats (500–700 g) were used in this study. All research protocols were approved and monitored by the Brown University Institutional Animal Care and Use Committee (IACUC), and all research was performed in accordance with relevant NIH guidelines and regulations. Rats were anesthetized with 2–3% isoflurane throughout the surgery. After shaving the hair, the animal was fixed on a stereotaxic frame, and the head skin was sterilized. Up to 5 cm sagittal incision was made in the skin over the skull, and a bilateral craniotomy was performed using a precision surgery dental drill. After the craniotomy, the skull in the midline was thinned down to improve the contact of neurograin interface. Insertion of neurograin electrodes was done manually and the array was fixed onto the skull with dental cement if needed. For interoperative recording and stimulation measurements, isoflurane was switched to 80 mg/kg Ketamine and 5 mg/kg Xylazine mixture and the depth of anesthesia was monitored every 5 minutes by checking the breath rate and paw retraction responses. If needed, ¼ of the initial dosage was given to the animal to maintain a stable anesthesia level. A heating pad was placed below the animal to maintain the body temperature, and the saline was administrated on the cortex occasionally to prevent drying of tissue. Upon completion of the surgery, animals were euthanized with pentobarbital sodium.

## Author contributions

J.L. designed RF wireless powering and data communication method, performed the system characterization, in vivo experiments, and analyzed the results. A.N. conceived the project with P.A., P.M. and L.L.. V.L., P.A., S.S. and L.L. designed RF data communication protocol. V.L., J.H., F.L., and P.M. designed the Neurograin ASICs. J.L. and A.N. designed the neural experimental concept. A.-H.L. performed microfabrication and packaging. J.L., V.L., F.L. and A.N. wrote the paper. All authors commented on the paper.

## Acknowledgements

The authors express their gratitude to Yoon-Kyu Song for insights to microfabrication and ASIC design, Joonsoo Jeong for his expertise with neurograin hermetic sealing and materials processing, Ramesh Rao for guidance on wireless networking design. We also thank Chester Kilfoyle, Ethan Mok, Stefan Sigurdsson, the Animal Care facility at Brown University for their most important contributions. We also acknowledge Siwei Li, Siyuan Yu, Lingxiao Cui, Sravya Alluri, Mustafa Lokhandwala at UCSD for their crucial work on ASIC design and benchtop test. The authors have benefited greatly from the insight of colleagues across multiple fields from microelectronics to brain sciences to clinical neurology: John Donoghue, Barundeb Dutta, John Groe, Leigh Hochberg, Terry Sejnowsky, Krishna Shenoy, Sydney Cash, among others. This research was initially supported by Defense Advanced Research Projects Agency N66001-17-C-4013, with subsequent support from private gifts. The postprocessing work benefited from equipment funded by the MRI award DMR-1827453.

