## Supplementary File for "Wireless Ensembles of Sub-mm Microimplants Communicating as a Network near 1 GHz in a Neural Application"

### Comparison with other contemporary approaches

Table S1 summarizes the performance of salient parameters of the neurograin approach and compares those with state-of-the-art wireless microimplants in terms of wireless powering, data communication, recording and stimulation. The Neurograin system is shown to have a considerably higher data rate to provide sufficient bandwidth for multi-node communication. Note that with the SoC design which integrates all circuit components as well as a coil on-chip, the present volume of Neurograin is approximately 0.1 $mm^3$, considerably smaller than other microdevices to date. We have elsewhere demonstrated that this volume can be reduced to 0.01 $mm^3$ after mechanical thinning and encapsulation with conformal hermetic packaging based on atomic-layer deposition (ALD) [1]. In addition to their unique capability for wireless networking, the miniaturized chips achieve recording and stimulation capability comparable to other microimplants. Designed for ultralow-power operation, the neurograin power budget is less than 30 μW for each chip. The main elements of the neurograin ASICs are summarized in Fig. S1.

**Table S1| Comparison between Neurograin and state-of-the-art microimplants**

| | [2] | [3] | [4] | [5] | [6] | [7] | Current work |
|---|---|---|---|---|---|---|---|
| Number of node(s) | 1 | 1 | 1 | 1 | 1 | 1 | 64 (up to 1000) |
| Wireless Powering | 1.85 MHz Ultrasonic | 1400 MHz RF Mid-field | 131 MHz RF 3-coil | 1180 MHz RF 2-coil | 433 MHz RF 3-coil | 1.85 MHz Ultrasonic | 915 MHz RF 3-coil |
| External Tx power (mW) | 0.12 | 800 | - | 4000 | - | 19.4 | Up to 500 |
| Operation depth (mm) | 8.8 (in-vivo) | 40 | - | 4.6 (beef) | - | 55 (ex-vivo) | 5-8 (phantom) |
| Telemetry | Backscattering | - | IR-UWB | - | Backscattering LSK | Backscattering | Backscattering BPSK (TDMA) |
| Uplink data rate (Mbps) | 0.5 | - | 0.8 | - | 0.205 | - | 10 |
| Type | Recording | Stimulation | Recording | Stimulation | Recording | Stimulation | Recording/ Stimulation |
| **Microimplants** | | | | | | | |
| Technology | Discrete | Discrete | 350 nm CMOS+ discrete | 130 nm CMOS | 350 nm CMOS | 65 nm CMOS | 65 nm CMOS |
| IC Area [$mm^2$] | 0.032 | - | 1.1 | 0.04 | 12.25 | 1 | 0.42 |
| Total Volume ($mm^3$) | 2.4 | 12 | 1 | 0.009 | 12 | 2.2 | 0.01-0.1 [1] |
| Energy harvester | Piezoelectric | Discrete coil | Discrete coil | On-chip coil | Discrete coil | Piezoelectric | On-chip Coil |
| Power supply (V) | - | - | 1.8 | 1.2 | 1.5 | 3 | 0.6-1 |
| Power Consumption (uW) | <1 | 500 | <300 | 8-38 | 92 | 65 | <30 |
| **Recording** | | | | | | | |
| Recorded signal | EMG | - | AP/LFP | - | LFP | - | LFP |
| Noise floor ($\mu V_{rms}$) | 180 | - | 3.78 | - | 1.8 | - | 2.2 |
| Resolution (bits) | 8 | - | 10 | - | 11 | - | 8 |
| Bandwidth (kHz) | > 30 | - | 10 | - | 0.825 | - | 0.5 |
| **Stimulation** | | | | | | | |
| Max. current (μA) | - | - | - | 46 | - | 400 | 10-25 |

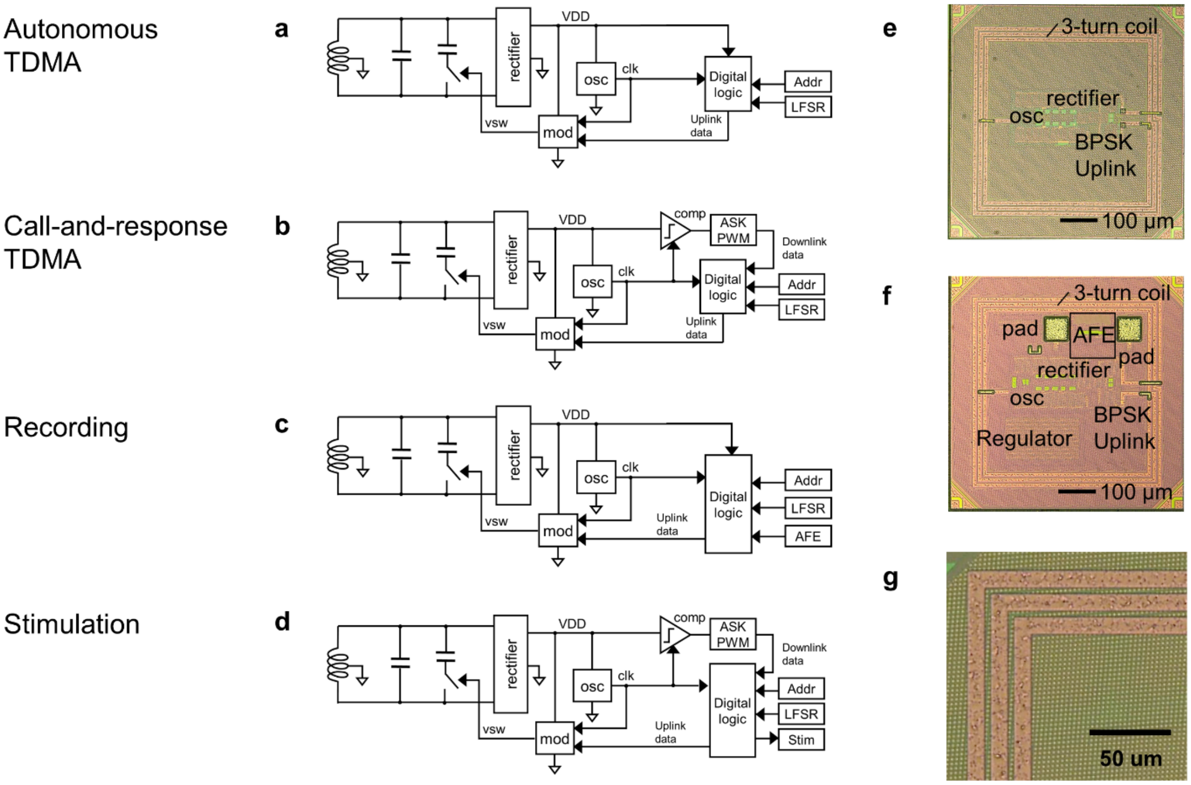


**Supplementary Figure 1| Circuit block diagrams for the Neurograin ASICs.** (a) Autonomous TDMA networking chip (with PUF address). Using a BPSK modulator, each neurograin backscatters every 100 ms while its data packet length is 100 us. The data packets contain about 1000 predefined bits from a linear-feedback shift register (LFSR) for BER performance evaluation. (b) Call-and-response TDMA chip, which shares RF energy harversting circuits with the autonomous version, but adds an amplitude shift keying, pulse width modulation (ASK-PWM) demodulator for bidirectional communication. (c) Recording ASIC with a PUF address and a capacitor-less analog-front-end (AFE). The recording chips have a voltage regulator to ensure stable recording. The chip backscatters in autonomous TDMA mode every 100 ms with the length of each data packet being 100 us. A packet contains predefined 32 bits from the LFSR, 20 bits dedicated to the PUF address, and 8000 bits for the ADC output (8 bits/100 samples) (d) Stimulating Neurograin with an ASK-PWM demodulator and a biphasic current source stimulator. The chip generates current output for injection to neural tissue when the address code in incoming downlink matches a 10-bit laser-written fuse address on-chip. Microphotographs of (e) The networking test chip, (f) A recording neurograin, (g) Spatial distribution of the top metal fill in the TSMC 65 nm process.

Abbreviations in the circuit diagrams. LFSR: linear-feedback shift register, Addr: address, mod: modulator, osc: oscillator, vsw: switch voltage, comp: comparator, AFE: analog-front-end, stim: stimulator

**Qualitative Description of the Neurograin Network**

The multi-node TDMA telecommunication between the external hub and an ensemble of spatially distributed implants is subject to overall latency. Our current networking protocol is designed for a 100 ms maximum latency, the latter serving as a constraint to the design of neurograin networking protocol. For instance, each recording neurograin simultaneously collects neural signals at 8 kbps (with ADCs sampling at 8-bit resolution and 1 kHz rate); 800 bits of data are thus accumulated over 100 ms per chip. To aid the data recovery at the external hub, data is packetized at each node with added frame structures such as pilot tones, PUF chip-IDs, cyclic-redundancy code (CRC), extending the overall packet size to 1000 bits.

To enable communication between a high number of nodes (e.g. 1000) and a single external RF hub, we adopted a TDMA (time-domain multiple-access) protocol. Each node, with its uniquely assigned identifier (ascending chip ID numbers), takes its turn to transmit data as in a daisy-chain configuration. For scaling up to a 1000-node wireless microimplant network, each node is given a small time window of 100 us (100 ms/ 1000 nodes) to transmit a data burst, thus collectively forming a 10 Mbps (1000 bit/ 100 us) uplink channel. To enhance network flexibility, we introduced a "call-and-respond" approach. The Chip-ID is embedded into the downlink trigger command, causing only the implant with the matching ID to report its data content. With the 16-bit PUF addresses, downlink rate of 1 Mbps, and Manchester-coding to ensure a data stream, each downlink trigger frame takes 40 us. Therefore, within the 100 ms latency window, 714 recording nodes can currently be addressed. Note that the call-and-response approach provides for a way to downselect chip nodes which may contain the most meaningful neural information, paving a way to use this type of bidirectional communication approach for future 'adaptive sensing' to reduce the burden on data processing and to further reduce the system latency.

Work is on-going to form an "adaptive daisy-chain" with close to 1000 nodes by implementing a "network schedule register" on the Neurograin. The idea is that ascending chip IDs are assigned (downloaded through the downlink) to an ensemble of neurograins selectively and prior to operation (calibration stage). The uplink could then be triggered with a much shorter downlink sequence during operation. As addresses only need to be downloaded occasionally, the channel can be utilized to almost 100% for recording/ stimulation data communication. The key concepts in our multi-node networking designs are summarized in Fig. S2, with experimental assessment in Figs. S3-S5.

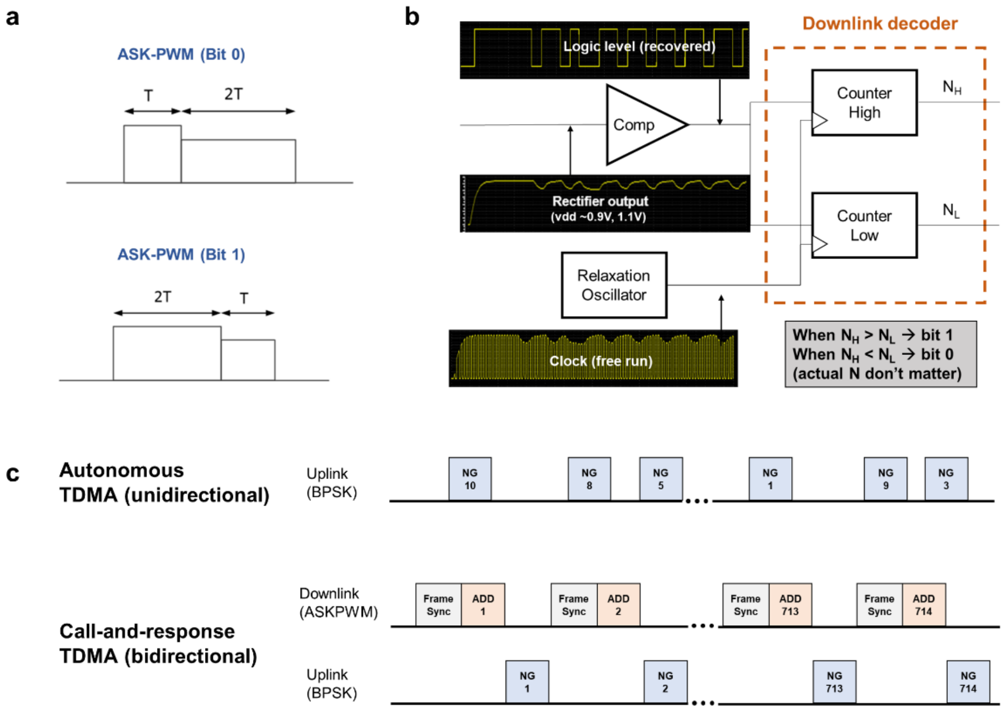


**Supplementary Figure 2| ASK-PWM scheme and networking protocol.** (a) ASK-PWM bits for data and frame synchronization; a short high represents bit 0 while a longer high is bit 1. (b) The logic decoder for relative pulse width comparison between high and low. (c) The timing diagram for autonomous and call-and-response TDMA. Both have a 100 ms frame and 100 us data packet length.

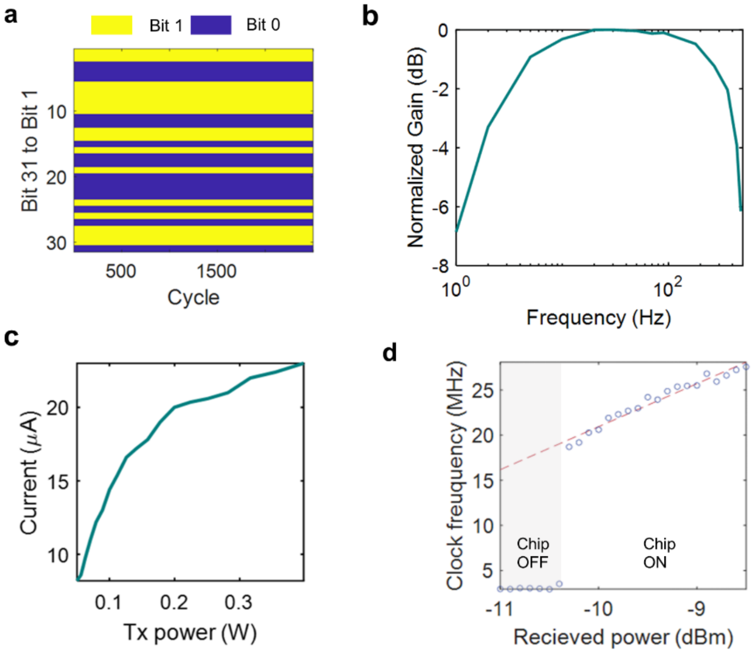


**Supplementary Figure 3| Results for ASIC test on analog-front end and data communication.** (a) Continuous BPSK demodulation results from a test chip with repeating 31 LFSR bits over 2500 cycles showing zero bit error. (b) Normalized gain spectrum of a recording neurograin; the low cutoff frequency is determined by the decoupling input capacitors in the test bench which mimics the high DC impedance in the electrode-electrolyte interface. (c) Injected current as a function of the external Tx power on a 2-coil test setup in a stimulating neurograin. (d) The relationship between the received power on the Rx coil and the clock frequency of the relaxation oscillator on-chip.

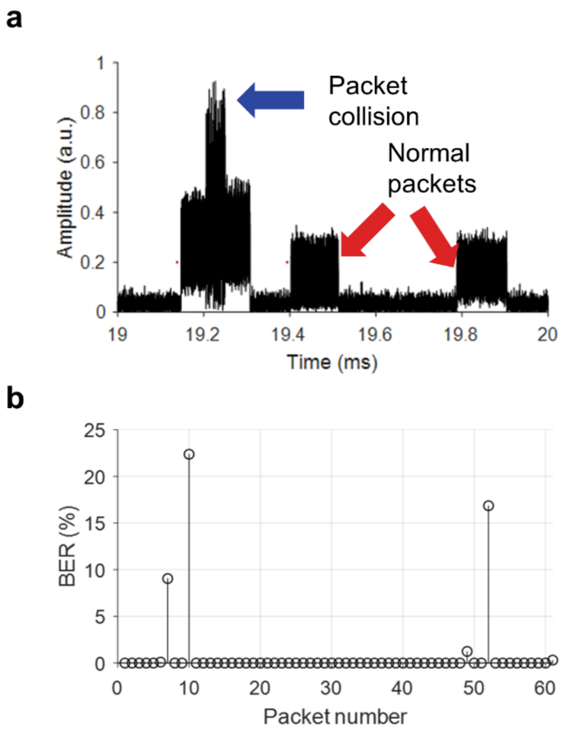


**Supplementary Figure 4| Autonomous TDMA demonstration.** (a) Transient waveform of received data showing uplink data packet collision from two neurograins in the autonomous TDMA mode. Collisions are caused by unscheduled backscattering and chip clock frequency variations. (b) Bit-error-rate (BER) of 61 packets measured over 100 ms from 64 autonomous TDMA chips. 4 packets show more than 1 % BER due to packet collision while the others have averaged BER of 0.007%.

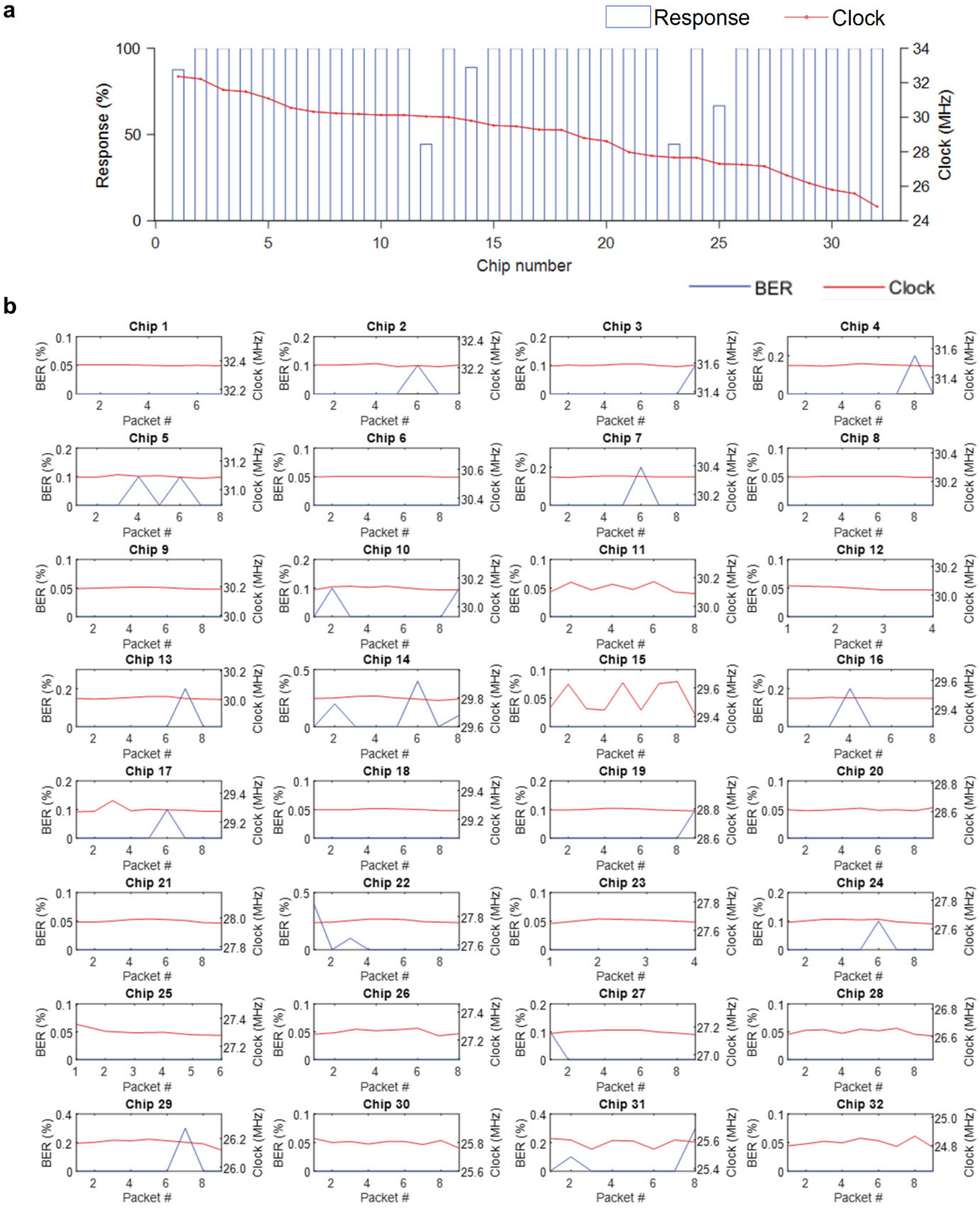


**Supplementary Figure 5| Measurement on call-and-response TDMA.** (a) ASK-PWM downlink response rate and the averaged clock frequencies of 32 call-and-respond TDMA chips. (b) Variations of BER and clock frequency from 9 received packets over the 32-chip network.

**Measured PUF performances on free-floating chips**

Shown in Fig. S6, in the case of an ensemble of 48 recording neurograins, 1697 data packets were detected during 3.3 seconds. Each packet contains a 20-bit PUF address, and exactly 48 distinct addresses were identified as in Fig. S6b. Using this information, we can uniquely associate each packet to the chip which sends it. Fig. S6a shows the counts of received packets per chip. Data packet collisions and clock speed differences contribute to the variation of packet counts. Fig. S6c shows the histogram of the number of different PUF bits between any two chips. It shows a normal distribution, with an average of 10 bits (that is, on average, half of the 20 PUF bits between two chips are different). Based on these observations, we conclude that the 20-bit PUF is random across chips, yet stable over time, thus making it a very suitable technique for chip addressing purposes for free-floating, ultra-small micro-implants.

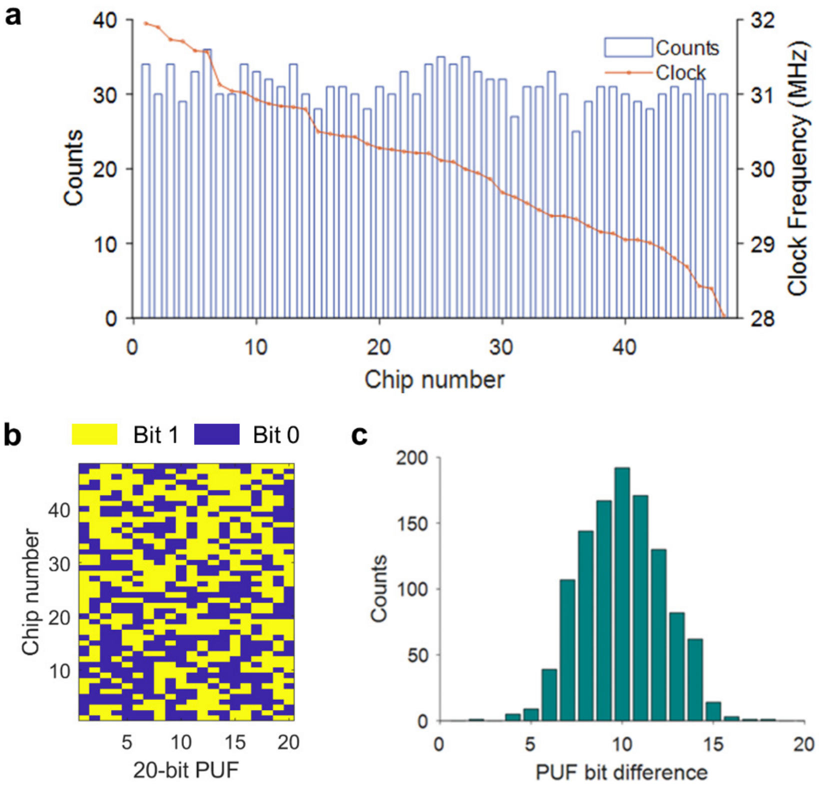


**Supplementary Figure 6| PUF analysis on 48 recording neurograin chips.** (a) Packet counts and clock frequencies, sorted using distinctive PUF addresses, of 48 chips measured over 3.3s. (b) The PUF addresses, each 20-bit long, of 48 chips. (c) The histogram on the number of bit differences (between two PUF addresses) for all 48 chips.

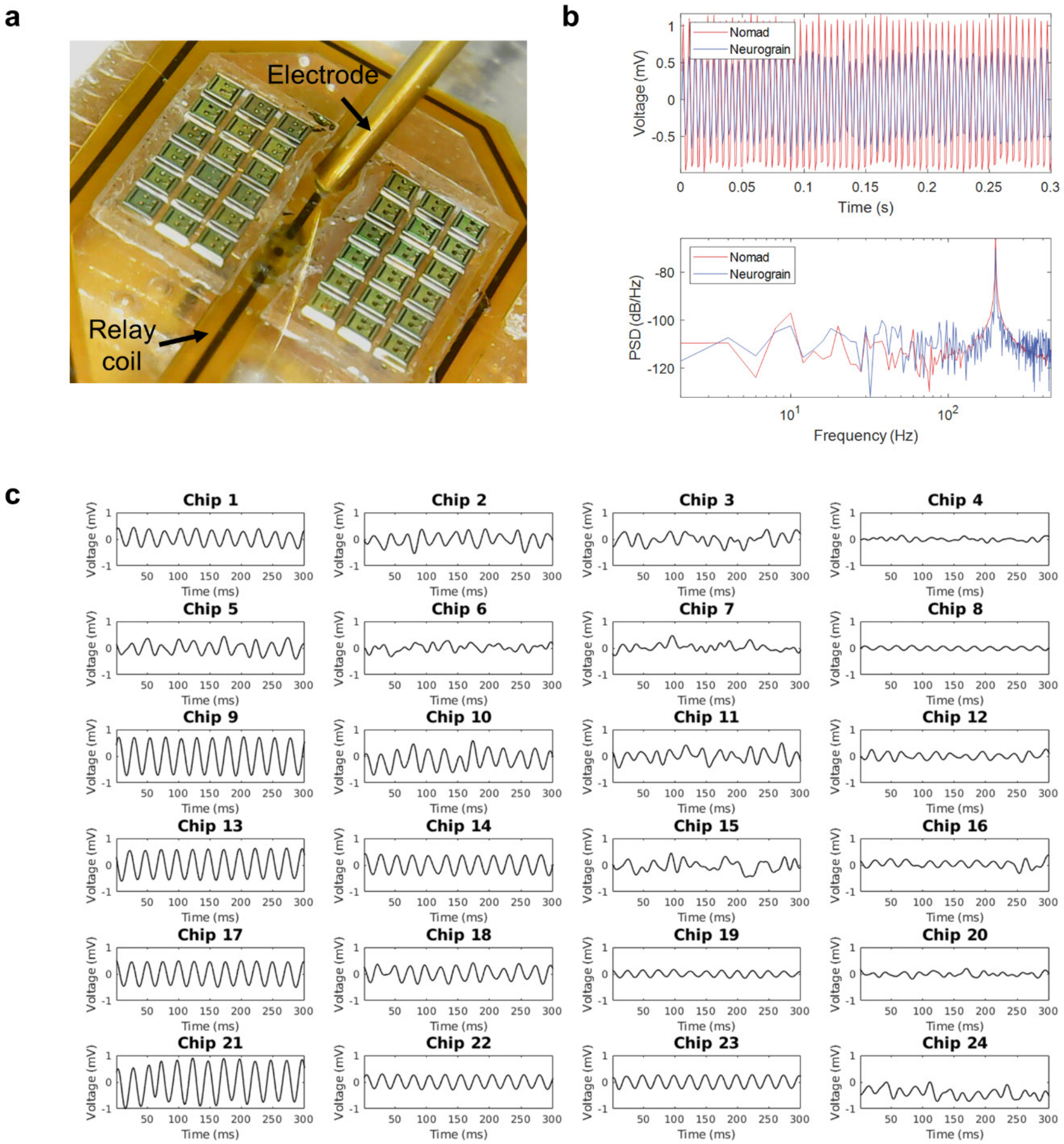


**Supplementary Figure 7| Recording neurograin test in saline.** (a) The photograph of the neurograin array in a saline testbench. The Tx and relay coil are underneath the array for wireless powering and data communication. (b) Recording of a 200 Hz signal injected in the saline by neurograin and a wired commercial device (Nomad, Ripple Inc) using gold planar electrodes. Both the transient waveforms (top) and the nonparametric power spectral density (PSD) estimates (bottom) are shown. (c) 40 Hz sinusoidal wave recordings by the distributed neurograin network in the saline (only 24 locations are shown).

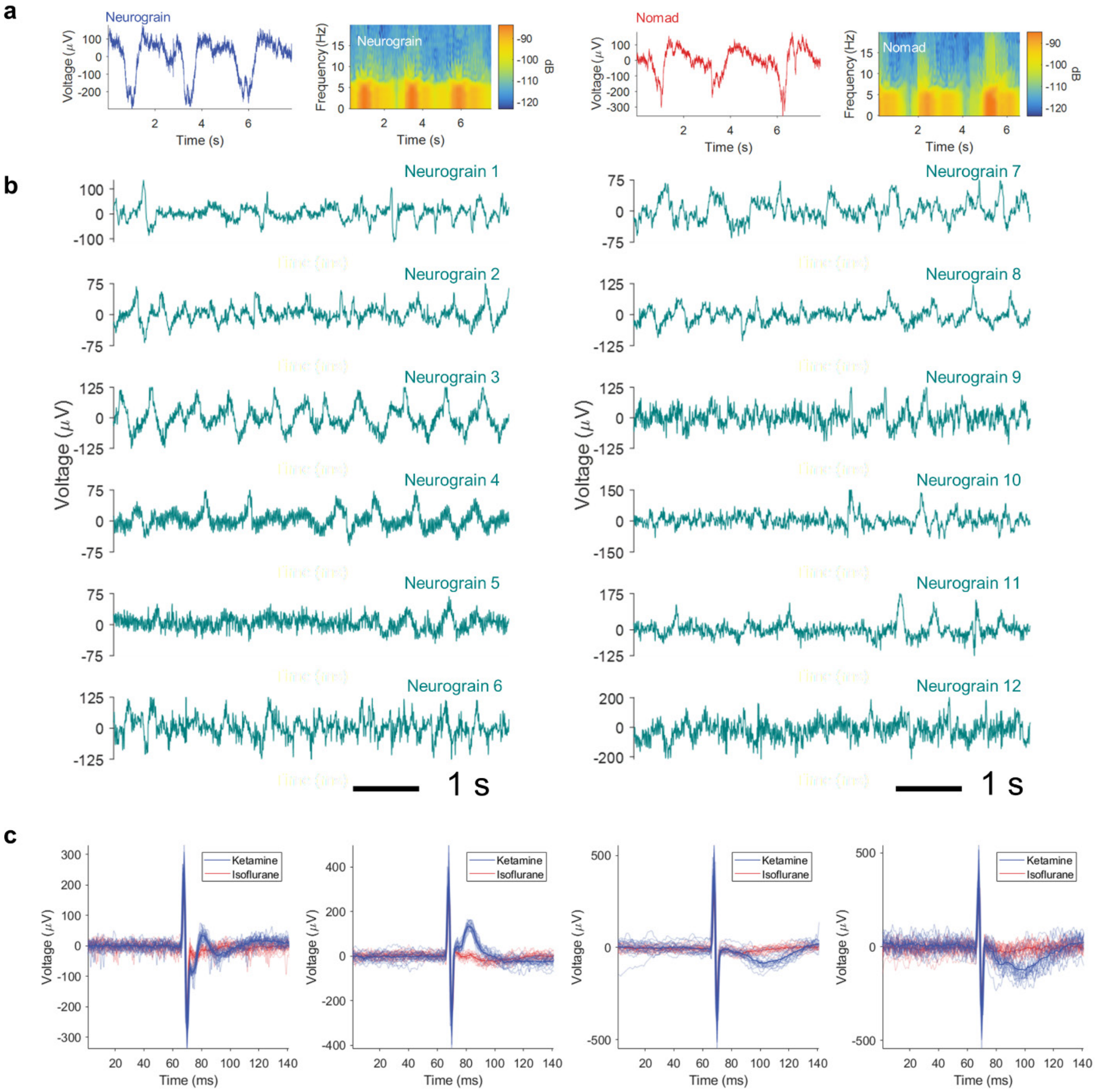


**Supplementary Figure 8| Demonstration of Neurograin recording in vivo**. (a) Neurograin and Nomad recordings of a spontaneous low-frequency oscillatory wave from anesthetized rat and their multi-taper spectrograms [8]. (b) Multiple neurograin recordings of spontaneous ECoG under ketamine which feature low-frequency oscillations. (c) Superimposed raw ECoG signals over 20 stimulation trials showing evoked responses with 20 to 70 milliseconds duration. The solid line shows averaged signals and each subplot shows results from different neurograins. The electrical stimulation has the amplitude of 50 uA and pulse width per phase is 1 ms.

**a**

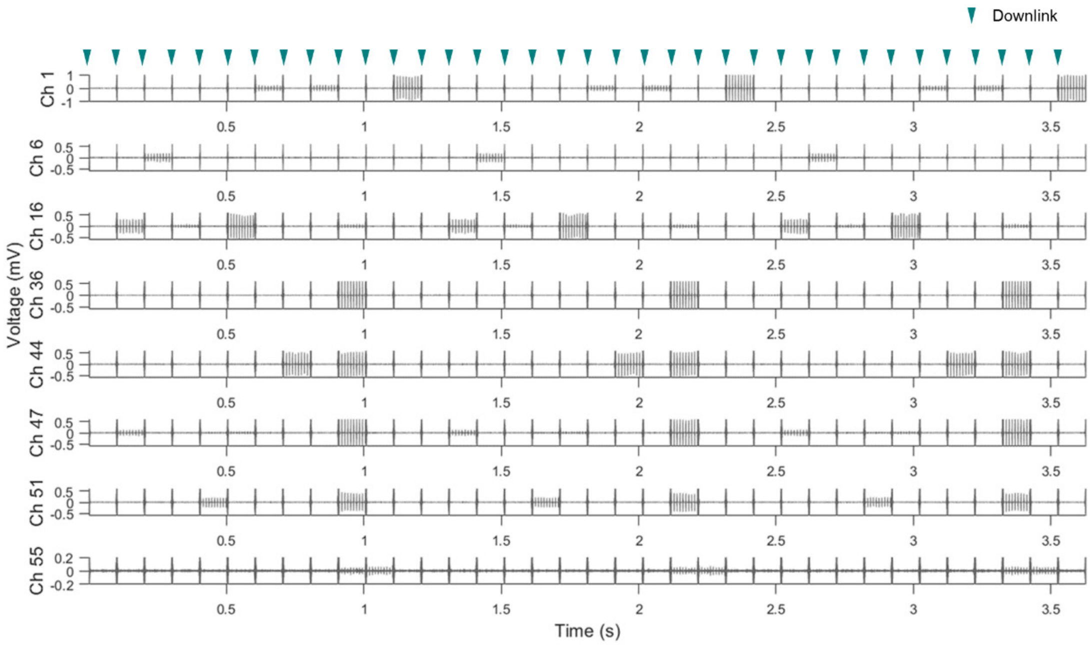


**b**

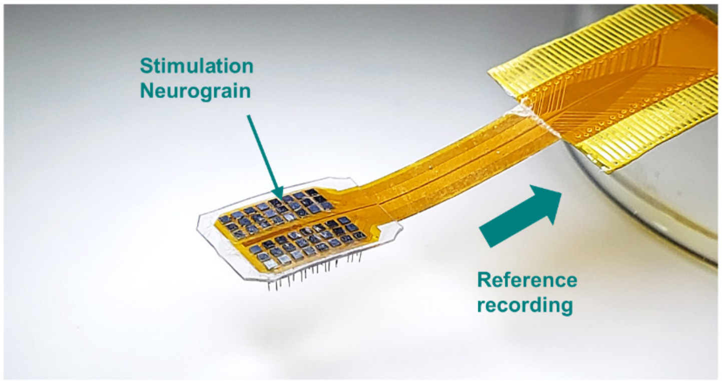


**Supplementary Figure 9| Stimulation Neurograin and stimulation patterns in saline.** (a) A stimulation pattern from 12 Neurograins recorded by 8 channels of a wired commercial device in saline. It shows the wireless programming (indicated by downlink arrows) of stimulation patterns over 3 cycles. (b) Stimulation neurograin array with post-processed penetrating electrodes and reference wired electrodes for monitoring. In this assembly, neurograins operate in a fully wireless manner and the reference electrodes in proximity monitor stimulation patterns and local neural activities to quantitively assess the effects of neurograin stimulation on the cortex.

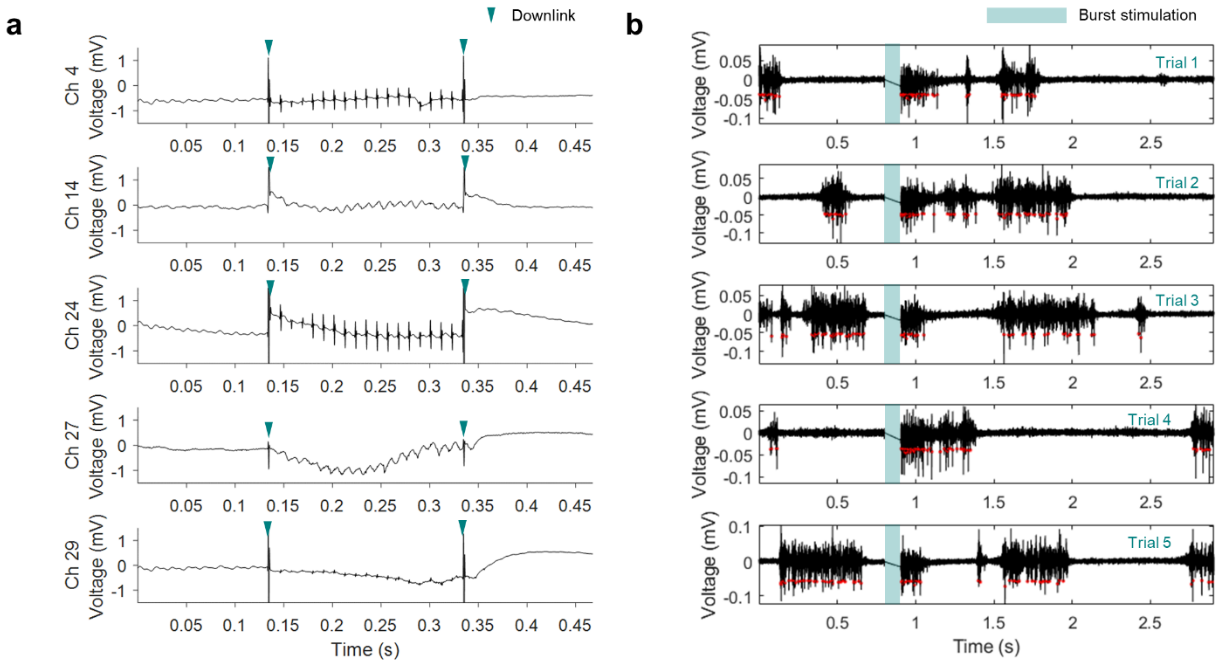


**Supplementary Figure 10| Cortical responses evoked by stimulation from a neurograin.** (a) 100 Hz Neurograin stimulation and evoked LFP responses over 5 recording channels. (b) Spontaneous burst firing activities under ketamine artificially induced by a 400 Hz Neurograin stimulation for 100 ms and recorded over 5 trials.

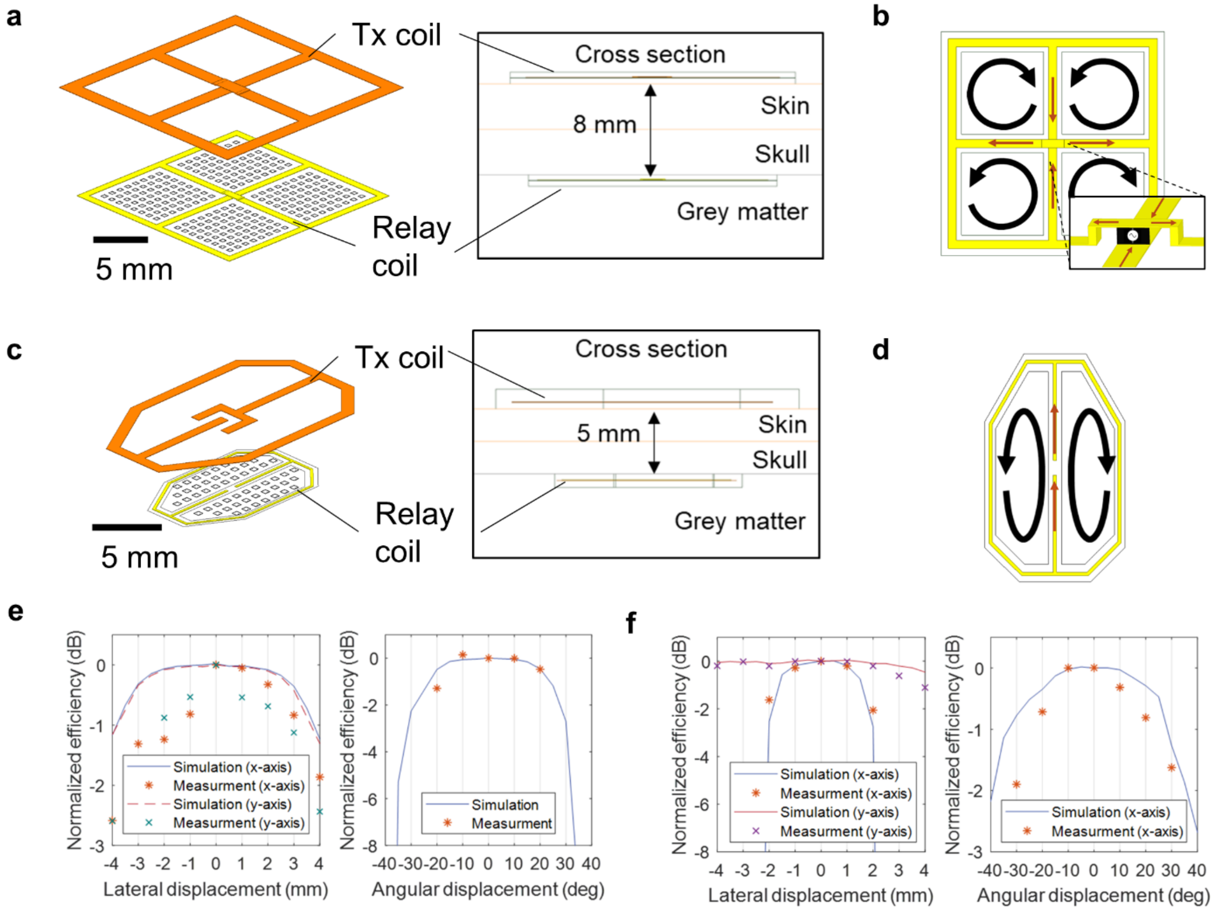


**Supplementary Figure 11| 3-Coil wireless power transfer system design and measurement.** (a) Wireless power transfer (WPT) system for the primate model consists of the transmitting (Tx) coil, the relay coil and the receiver (Rx) micro coils. (c) Coils for the rodent model. (b) & (d) Four or two sub-coils are combined to realize specific current flow for the primate and the rodent models, respectively. They expand the powering area for scalability. (e) & (f) WPT efficiency as a function of the lateral or angular displacement of Tx coil for the primate and the rodent models, respectively.

| Table S2\| Dimension and simulation parameters of WPT coil system at 915 MHz | | |
|---|---|---|
| **Type** | **Primate model coil** | **Rodent model coil** |
| $L_1$ (nH) | 12.2 | 20.9 |
| $L_R$ (nH) | 11 | 16 |
| $L_2$ (nH) | 10.3 | |
| $Q_1$ | 26.9 | 50 |
| $Q_R$ | 30.8 | 109.8 |
| $Q_2$ | 13.4 | |
| $k_{1R}$ | 0.051 | 0.02 |
| $k_{R2}$ | 0.0035-0.0099 | 0.008-0.015 |
| $k_{12}$ | 0.0005-0.0006 | 0.0003-0.0007 |
| Size Tx (mm) | 22.5 x 22.5 | 20 x 12.3 |
| Size Relay (mm) | 20.4 x 20.4 | 14.2 x 8 |
| Size Rx (mm) | 0.5 x 0.5 (3 turns) | |
| Tissue | 4 mm skin, 4 mm skull, >20 mm cortex | 2.5 mm skin, 2.5 mm skull, >20 mm cortex |

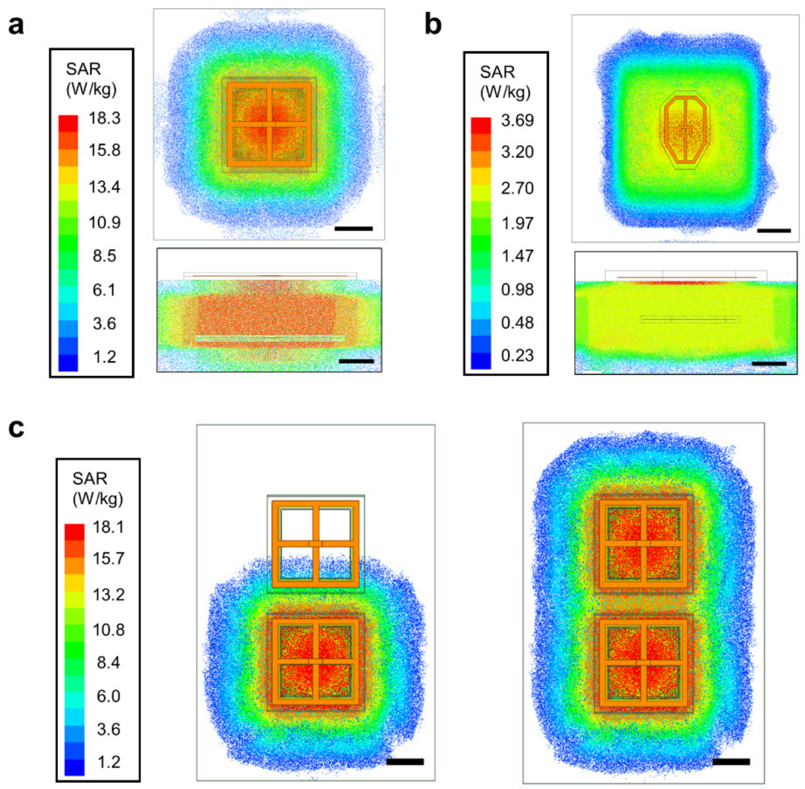


**Supplementary Figure 12| Specific absorption rate (SAR) simulation.** (a) SAR field pattern of the primate WPT system with 0.5 W Tx emission. (b) SAR field of the rodent model coil with 0.072 W Tx emission. (c) Comparison of SAR fields with one Tx channel enabled (left) and two adjacent Tx channels enabled (right) with 0.5 W emission for each. The scale bar is 10 mm.

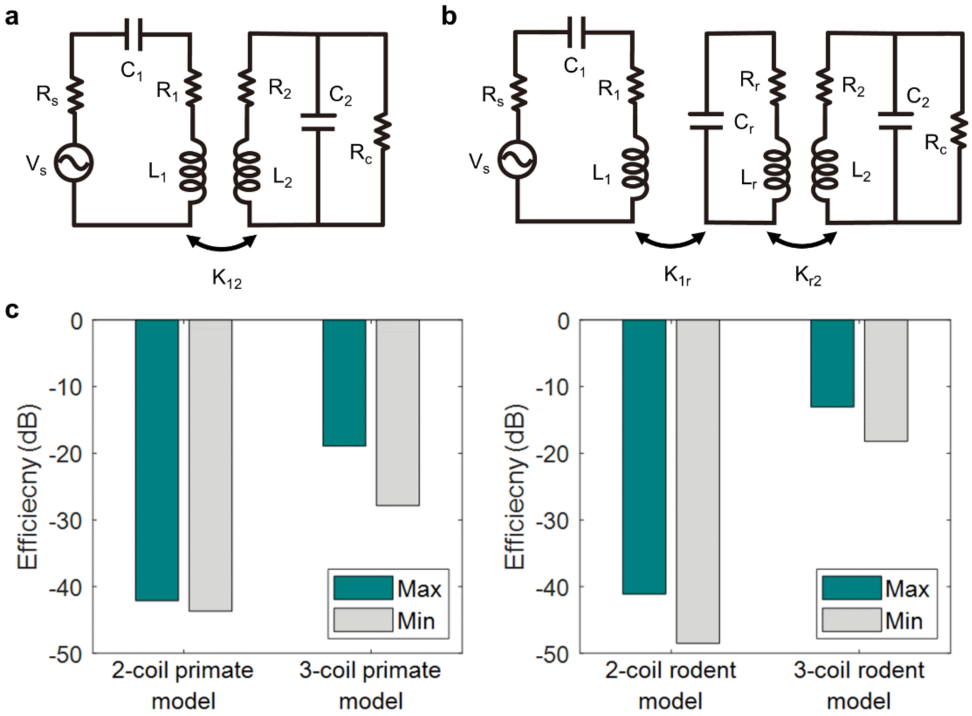


**Supplementary Figure 13| Comparing the 2-coil vs. 3-coil WPT systems.** (a) The lumped circuit model for the 2-coil wireless powering system. (b) The 3-coil model. (c) 2-coil vs 3-coil efficiency comparison for the primate and rodent systems, using Equation 1 and 2 with coil parameters from Table S2. Vs: source voltage, Rs: source impedance, Rc: circuit impedance.

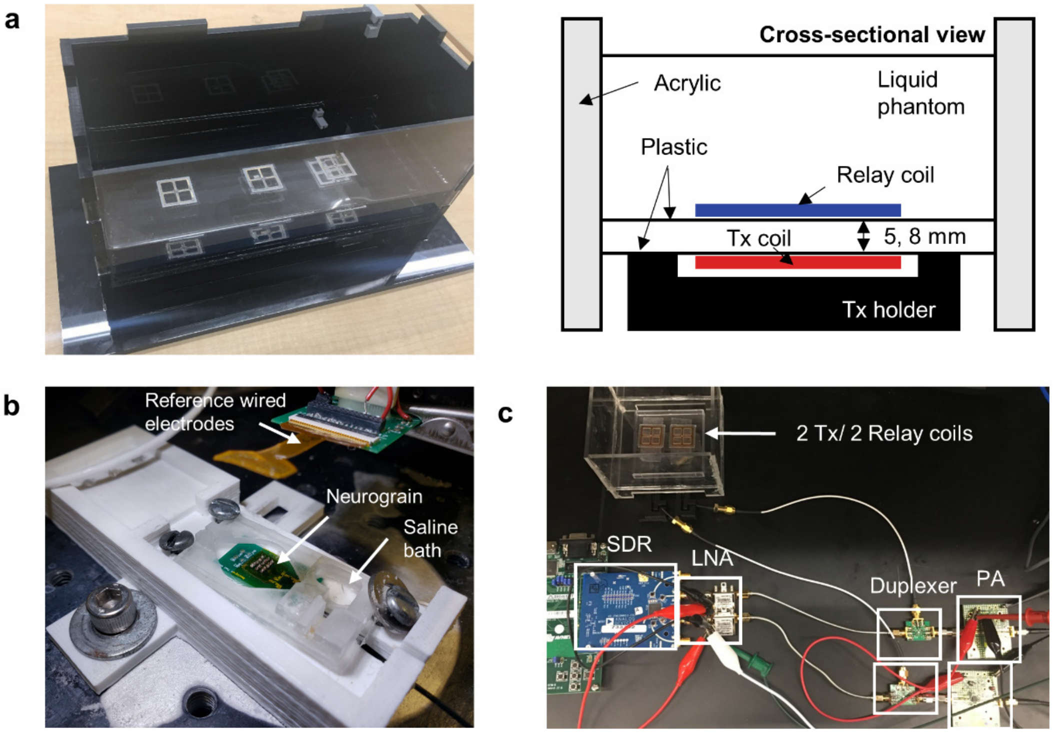


**Supplementary Figure 14| Neurograin benchtop RF setup.** (a) Acrylic and plastic-based setup to minimize RF interference, and a liquid phantom to mimic dielectric properties of brain tissues [9]. (b) Saline bath for testing recording/stimulation neurograins along with reference wired electrodes. (c) RF set up for dual channel power and data link using two adjacent pairs of Tx/ relay coils to double the channel capacity.
